# NHE1 sustains leukemia-propagating capacity and mitochondrial fitness in T-cell acute lymphoblastic leukemia

**DOI:** 10.64898/2026.10.18.759977

**Authors:** Majd A. Al-Hamaly, Meghan G. Haney, Yelena Chernyavskaya, Ty C. Cheatham, Jeffery T. Jolly, Michelle G. Pitts, Daheng He, Logan T. Turner, Ronald C. Bruntz, Chi Wang, Jessica S. Blackburn

## Abstract

Leukemia-propagating cells (LPCs) sustain T-cell acute lymphoblastic leukemia (T-ALL) and are thought to contribute to disease progression and relapse. However, the lack of reliable LPC markers has hindered efforts to identify their vulnerabilities. We screened 786 FDA-approved compounds using a zebrafish transplantation assay that directly measures leukemia-propagating activity. Amiloride markedly reduced functional LPC frequency (*p* < 0.0001) without significantly delaying leukemia onset at higher transplantation doses. Because amiloride inhibits the sodium-hydrogen exchanger NHE1, we investigated its role in T-ALL. NHE1 knockdown reduced self-renewal-associated colony formation by >70% human T-ALL cells, with only modest increases in apoptosis. Transcriptomic and proteomic analyses revealed extensive molecular remodeling, including reduced abundance of proteins involved in mitochondrial respiration and translation. Amiloride treatment and NHE1 knockdown impaired mitochondrial respiration and reduced mitochondrial content, while NHE1 knockdown also decreased mitochondrial membrane potential. Metabolomic profiling revealed alterations in TCA cycle intermediates and broader cellular metabolism. In mouse xenografts, NHE1 knockdown reduced leukemia involvement in the bone marrow and spleen despite similar overall leukemia progression and survival. Together, these findings implicate NHE1 as a previously unrecognized regulator of T-ALL leukemia-propagating activity, reveal unexpected contributions of this ion exchanger to mitochondrial fitness and leukemia involvement in hematopoietic tissues, and support further investigation of NHE1 as a potential therapeutic target in T-ALL.

**Key Points:**

- NHE1 sustains leukemia propagating capacity and self-renewal in T-ALL
- NHE1 knockdown impairs mitochondrial fitness and reduces leukemia involvement in hematopoietic tissues.

## Introduction

T-cell acute lymphoblastic leukemia (T-ALL) is an aggressive hematologic malignancy for which multi-agent chemotherapy has substantially improved outcomes. Despite these advances, approximately 10–15% of children experience relapse, and outcomes after relapse remain poor across age groups, with long-term survival often below 30% (1,2). Disease persistence and relapse are thought to be driven by leukemia cells with the capacity to self-renew and regenerate leukemia, yet these leukemia-propagating cells are not reliably defined by a single immunophenotypic population in T-ALL (3,4). This heterogeneity has made it difficult to identify vulnerabilities that specifically regulate self-renewal and has limited the development of therapies that target the leukemia-propagating compartment. Functional approaches that directly measure self-renewal and leukemia-propagating capacity therefore provide an opportunity to uncover therapeutic dependencies that may be missed by marker– or target-based strategies.

To address this challenge, we used leukemia clones enriched for leukemia-propagating cells (∼10%), derived from the transgenic *rag2-Myc* zebrafish T-ALL model(5,6). This unusually high frequency of leukemia-propagating cells enables self-renewal to be interrogated functionally in vivo using transplantation-based assays. We therefore used these clones to perform a phenotypic, large-scale screen of 786 FDA-approved compounds for their ability to suppress leukemia-propagating activity. By beginning with an in vivo functional screen and then advancing candidate compounds through complementary assays in human T-ALL cells, we sought to identify druggable regulators of self-renewal without requiring a predefined molecular target.

This screening strategy identified amiloride as a top candidate inhibitor of T-ALL self-renewal. Amiloride is an orally administered potassium-sparing diuretic that, in addition to its effects on epithelial sodium channels, inhibits the sodium-hydrogen exchanger-1 (NHE1; encoded by *SLC9A1*) (7,8). NHE1 exchanges intracellular H+ for extracellular Na+ and is a major regulator of intracellular pH and cellular ion homeostasis(9,10). Previous studies have implicated NHE1 in leukemia migration and therapeutic resistance, including in T-ALL(11–15). However, its role in T-ALL self-renewal has not been defined, and the mechanisms through which NHE1 might support leukemia-propagating cells remain unknown. We therefore sought to determine whether NHE1 regulates T-ALL self-renewal and to define the cellular pathways underlying this phenotype.

Here, we identify NHE1 as a previously unrecognized regulator of T-ALL self-renewal. Pharmacologic and genetic perturbation of NHE1 converged on impaired leukemia-propagating activity, while unbiased transcriptomic and proteomic profiling revealed mitochondrial dysfunction as a prominent consequence of NHE1 loss. Functional and metabolomic studies further demonstrated that NHE1 perturbation compromises mitochondrial bioenergetics and integrity and broadly disrupts cellular metabolism, while in vivo studies revealed reduced leukemia involvement in hematopoietic tissues.

These findings identify NHE1 as a regulator of T-ALL self-renewal and define mitochondrial, metabolic, and tissue-specific consequences of NHE1 loss.

## Methods

### Zebrafish drug screening and limiting-dilution transplantation

Previously characterized GFP-positive zebrafish T-ALL clones 1.3 and 10.2 were propagated in syngeneic CG1 recipients. Leukemia cells were incubated for 1 hour with pools of eight compounds from a 786-compound FDA-approved drug library (1 µM per compound) and transplanted intraperitoneally into 15 recipients per pool at 25 cells per fish. Leukemia development was monitored for 6 weeks. Fifty-six compounds from selected pools were individually evaluated in Jurkat colony-formation assays, followed by viability assessment of selected hits. For limiting-dilution transplantation, clone 1.3 cells were exposed to 1 µM compound or vehicle for 1 hour and transplanted at 10, 100, or 1,000 cells per recipient. Leukemia-propagating cell frequencies were estimated using extreme limiting dilution analysis (ELDA). Zebrafish studies were approved by the University of Kentucky Institutional Animal Care and Use Committee (IACUC).

### Human T-ALL cells and functional assays

Jurkat, CCRF-CEM, MOLT-4, and SUP-T1 cells were cultured in RPMI 1640 supplemented with 10% fetal bovine serum and routinely tested for mycoplasma. NHE1 was knocked down in Jurkat cells using lentiviral *SLC9A1*-targeting shRNAs and scrambled controls, with knockdown verified by immunoblotting and RT-qPCR. Cells were treated with amiloride, hexamethylene amiloride, or appropriate vehicle controls as indicated. Colony formation was quantified after 14 days in methylcellulose (2,000 cells/well). Apoptosis was assessed by Annexin V staining and cell-cycle distribution by propidium iodide DNA-content analysis.

### Mitochondrial function and molecular profiling

Oxygen consumption was measured using an Agilent Seahorse XF Pro analyzer with sequential oligomycin, FCCP, and rotenone/antimycin A injections. Basal respiration, ATP-linked respiration, and spare respiratory capacity were calculated. Mitochondrial membrane potential and content were evaluated using JC-1 and MitoView Green staining, respectively. Confocal images of MitoView Green– labeled cells were analyzed using ImageJ/MiNA to quantify mitochondrial footprint. RNA sequencing of NHE1-knockdown and control Jurkat cells was analyzed using STAR, HTSeq, DESeq2, and gene set enrichment analysis. Proteins were profiled using liquid chromatography–tandem mass spectrometry, with differential abundance and pathway analyses using DEP and WebGestalt. Metabolites were analyzed using gas chromatography–mass spectrometry and MetaboAnalyst.

### Analysis of TARGET-ALL-P2 data

TPM-normalized RNA-seq data from 463 primary and 69 partially paired recurrent ALL samples were obtained from the GDC Data Portal. *SLC9A1* expression was compared using a random-intercept linear mixed-effects model. Survival data from cBioPortal were analyzed using Kaplan–Meier curves, log-rank tests, and Cox proportional hazards regression. The high-versus low-expression cutoff was selected using *maxstat*, with adjustment of log-rank *p* values for cutoff optimization.

### Mouse xenografts and tissue analyses

Luciferase-expressing Jurkat cells carrying scrambled or NHE1-targeting shRNAs were injected intravenously into 6-to 8-week-old NSG mice at 10⁵ or 10⁶ cells per mouse (control, n = 6; knockdown, n = 7). Leukemia development was monitored by bioluminescence imaging for up to 10 weeks. Bone marrow leukemia burden was measured by human CD45 flow cytometry; apoptosis was evaluated by cleaved caspase-3 immunohistochemistry. Mouse studies were approved by the University of Kentucky IACUC.

### Statistical analyses

Comparisons were performed using t tests or one– or two-way ANOVA with appropriate multiple-comparison corrections, as specified in the figure legends. Sample sizes and statistical tests are reported in the corresponding figure legends. Additional experimental procedures are provided in the supplemental Methods.

## Results

### Sequential zebrafish and human T-ALL screening identifies candidate inhibitors of leukemia-propagating cell function

To identify compounds that functionally impair T-ALL self-renewal, we screened two previously established *rag2-Myc* zebrafish T-ALL clones (1.3 and 10.2) with leukemia-propagating cell (LPC) frequencies of approximately 10%(5). Because LPC frequency in these clones is approximately 1 in 10 cells, we transplanted 25 cells per recipient rather than saturating cell numbers, making leukemia establishment sensitive to compounds that impair LPC function. Leukemia cells were exposed ex vivo to pools of eight compounds and transplanted with drug into syngeneic recipients. In total, 786 compounds were evaluated in 3,060 recipient zebrafish across two T-ALL samples (**Figure 1A**).

**Figure 1.**
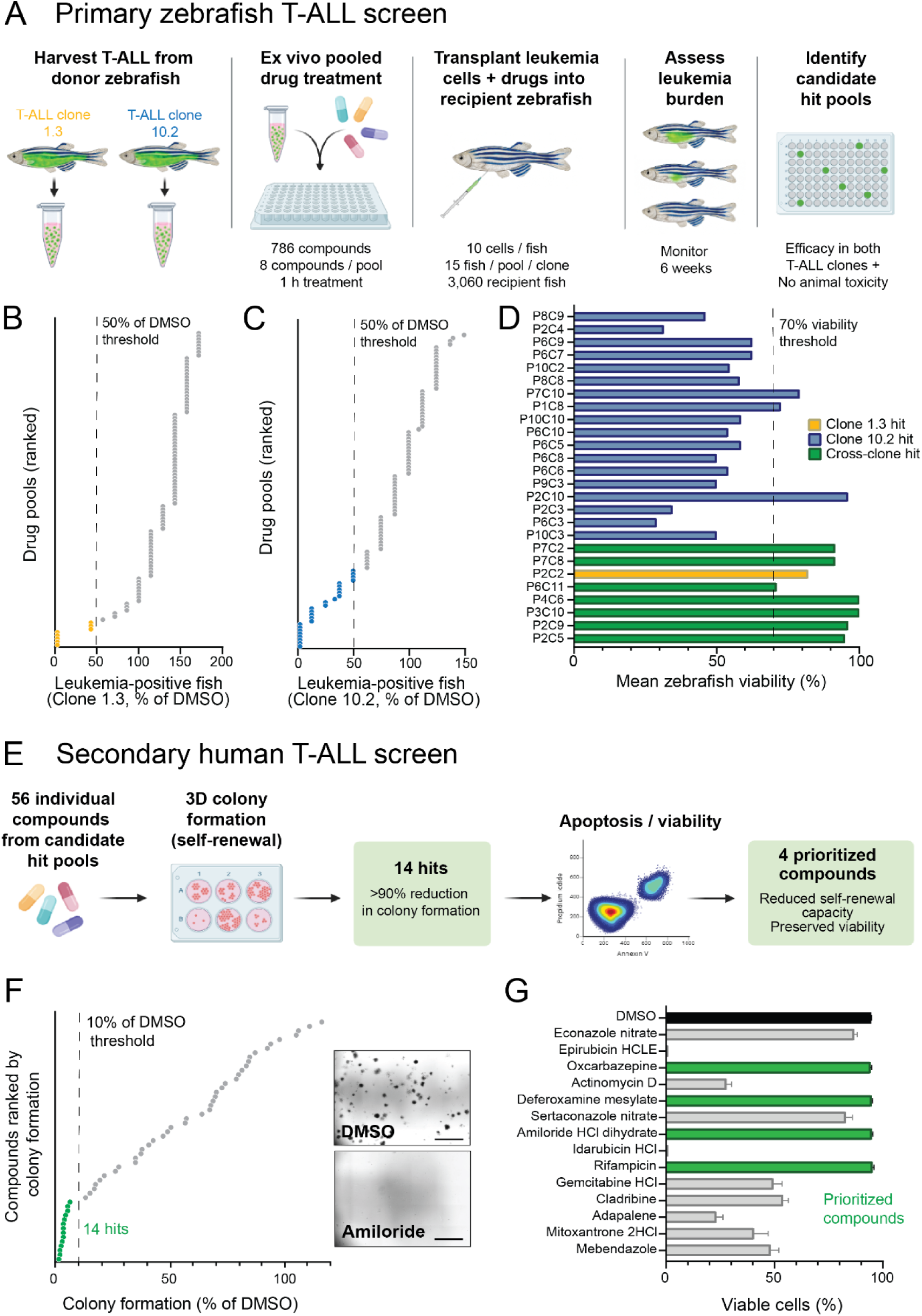
Sequential zebrafish and human T-ALL screening identifies compounds that suppress leukemia-propagating and self-renewal activity. (A) Schematic of the primary zebrafish drug screen. Leukemia cells from two independently derived *rag2-myc* T-ALL clones (1.3 and 10.2) were harvested from donor zebrafish, exposed ex vivo for 1 h to pools of 8 compounds from a 786-compound FDA-approved drug library, and transplanted with drug into syngeneic CG1 recipient zebrafish at 10 cells per fish (15 recipients per pool per clone; 3,060 recipient fish total). Leukemia development was assessed over 6 weeks. (B-C) Drug pools ranked by the percentage of leukemia-positive recipients relative to DMSO for clone 1.3 (B) and clone 10.2 (C). Dashed lines indicate 50% of the DMSO leukemia incidence. Gold and blue points indicate pools meeting this threshold in clones 1.3 and 10.2, respectively. (D) Mean recipient viability for candidate hit pools, ranked by anti-leukemia activity across the two clones. The dashed line indicates the 70% viability threshold. Gold bars indicate clone 1.3-specific hits, blue bars indicate clone 10.2-specific hits, and green bars indicate hits identified in both clones. (E) Schematic of the secondary human T-ALL screen. The 56 individual compounds contained within candidate hit pools were evaluated in a MethoCult colony-formation assay. Fourteen compounds that reduced colony formation by >90% were advanced to apoptosis/viability testing, yielding 4 prioritized compounds that retained >90% viability. (F) Colony formation following treatment with individual compounds (10 μM), expressed relative to DMSO and ranked by response. The dashed line indicates 10% of DMSO colony formation (>90% reduction); green points denote the 14 compounds meeting this criterion. Representative images of DMSO– and amiloride-treated colonies are shown at right. Scale bar, 2 mm. **(G)** Percentage of viable Jurkat cells following 24-h treatment with the 14 colony-formation hits (10 μM), assessed by Annexin V/PI staining. Green bars indicate the 4 prioritized compounds that retained >90% viability while strongly suppressing colony formation.

Drug-pool activity was quantified as the percentage of leukemia-positive zebrafish relative to DMSO in each T-ALL clone (**Figure 1B-C**). Twenty-six pools reduced leukemia incidence to ≤50% of DMSO in at least one clone and were further evaluated for effects on animal viability (**Figure 1D**). Clone 10.2 was more broadly drug responsive than clone 1.3; requiring activity across both independently derived leukemias provided a stringent filter for reproducible effects on leukemia propagation. Seven drug pools reduced leukemia incidence to ≤50% of DMSO in both clones while maintaining >70% mean zebrafish viability. These pools contained 56 individual compounds that were advanced to secondary screening. Complete in vivo primary-screen efficacy and viability data are provided in **Supplemental Table 1**.

To determine whether these compounds also impaired human T-ALL self-renewal capacity, all 56 compounds were tested individually at 10 μM in a Jurkat MethoCult colony-formation assay, used as a functional in vitro surrogate of self-renewal (**Figure 1E**) (16). Fourteen compounds reduced colony formation to ≤10% of DMSO after 14 days (**Figure 1F**; **Supplemental Table 1**). Because loss of colony formation could also result from nonspecific cell death, these 14 compounds were subsequently evaluated by Annexin V/PI staining after 24 hours of treatment. Four compounds, amiloride, deferoxamine, rifampicin, and oxcarbazepine, maintained >90% cell viability while reducing colony formation by >90% (**Figure 1G**, **Supplemental Figure 1**). These compounds were therefore advanced for direct evaluation of leukemia-propagating activity.

### Amiloride reduces leukemia-propagating cell frequency in vivo

We next tested whether the four candidates identified by our screening pipeline altered functional leukemia-propagating capacity *in vivo* using limiting-dilution transplantation. Zebrafish T-ALL clone 1.3 cells were treated with each compound or DMSO and transplanted at 1,000, 100, or 10 cells per recipient. Among the four candidates, amiloride produced the largest reduction in leukemia-propagating cell frequency, decreasing the estimated frequency from 1 in 10.2 cells in DMSO-treated controls to 1 in 149.6 cells following treatment (*p* < 0.0001), while oxcarbazepine produced a smaller but significant reduction (*p* = 0.004) and deferoxamine and rifampicin had no significant effect (**Figure 2A-B; Supplemental Table 2**). At the higher 100– and 1,000-cell doses, where leukemia developed in nearly all recipients, the time to leukemia onset was similar between DMSO– and amiloride-treated groups (**Figure 2C**; *p* = 0.151).

**Figure 2.**
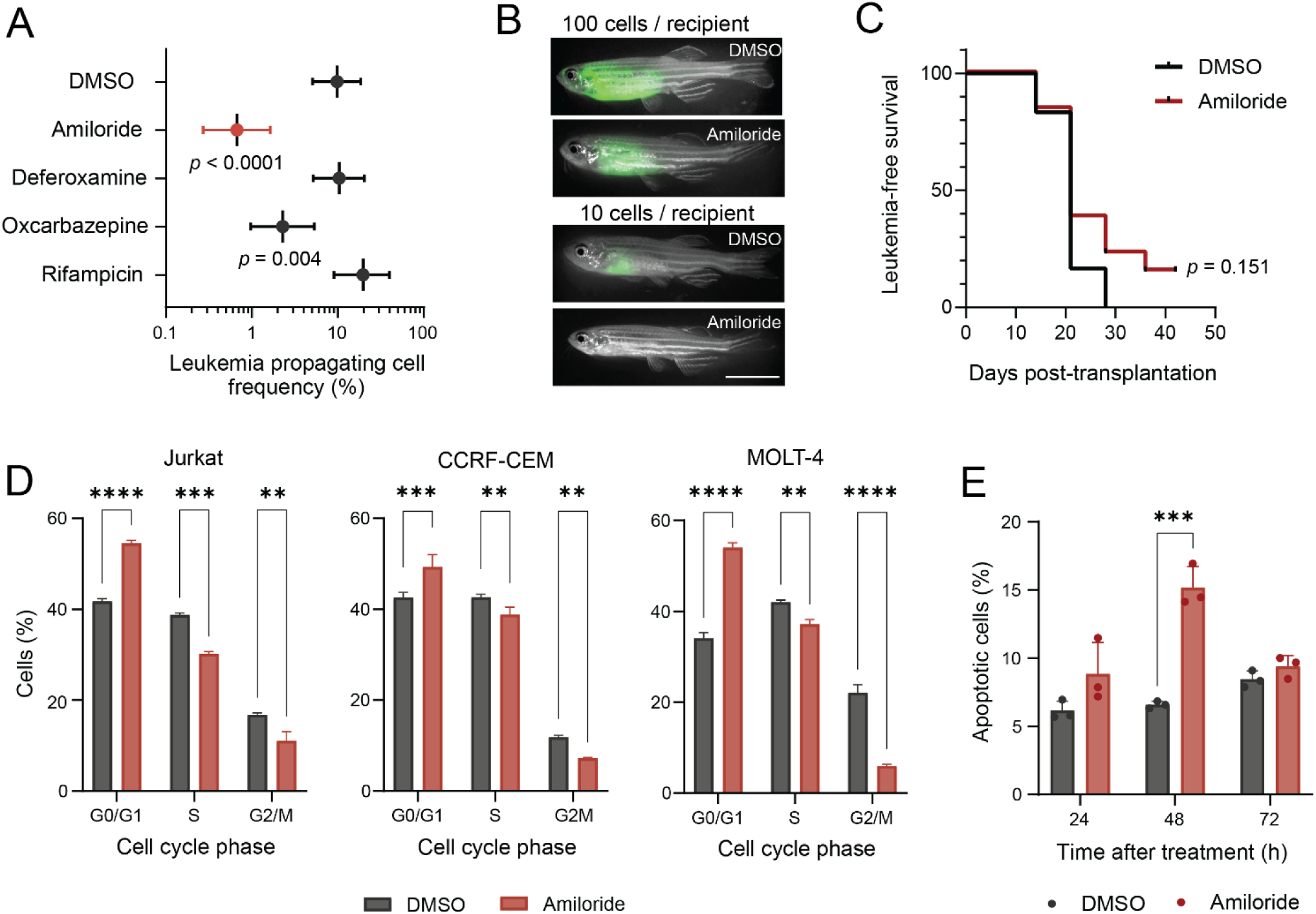
Amiloride reduces leukemia-propagating cell frequency in T-ALL. **(A)** Estimated leukemia-propagating cell (LPC) frequency following limiting-dilution transplantation of YFP-positive zebrafish T-ALL clone 1.3 treated with DMSO or the indicated compounds. Cells were transplanted at 1,000, 100, or 10 cells per recipient. Points indicate LPC frequency estimates with 95% confidence intervals determined by extreme limiting dilution analysis (ELDA) and plotted on a logarithmic scale. *p* values indicate comparisons with DMSO by ELDA; nonsignificant comparisons are not shown. **(B)** Representative recipients transplanted with 100 or 10 DMSO– or amiloride-treated T-ALL cells and imaged 28 days after transplantation. Scale bar, 10 mm. **(C)** Leukemia-free survival in the pooled 1,000– and 100-cell transplantation cohorts following treatment with DMSO or amiloride. *p* = 0.151 by log-rank (Mantel-Cox) test. **(D)** Cell-cycle distribution of Jurkat, CCRF-CEM, and MOLT-4 cells following 24-h treatment with DMSO or 100 μM amiloride, assessed by propidium iodide staining. Bars indicate the mean of three independent biological replicates ± SD (*n* = 3). **(E)** Total apoptotic Jurkat cells following treatment with DMSO or 100 μM amiloride for 24, 48, or 72 h, assessed by Annexin V/propidium iodide staining. Points represent independent biological replicates; bars indicate mean ± SD (*n* = 3). Statistical significance in **D-E** was determined by two-way repeated-measures ANOVA with Sidak’s multiple-comparisons correction. ** *p* < 0.01; *** *p* < 0.001; **** *p* < 0.0001.

In human T-ALL cells, amiloride treatment produced statistically significant but relatively modest changes in cell-cycle distribution across Jurkat, CCRF-CEM, and MOLT-4 cells, with an increased G₀/G₁ fraction and corresponding reductions in S and G₂/M populations (**Figure 2D; Supplemental Figure 2A**). Amiloride also induced a modest increase in apoptosis at 48 hours that was not sustained at 72 hours (**Figure 2E; Supplemental Figure 2B**). These data demonstrate that amiloride markedly impairs T-ALL self-renewal and leukemia-propagating activity without causing widespread cytotoxicity or a comparable suppression of bulk leukemia growth.

### NHE1 depletion suppresses self-renewal and alters cell cycle progression in T-ALL cells

Amiloride is a known inhibitor of the Na+/H+ exchanger 1 (NHE1), encoded by *SLC9A1*(17). We therefore asked whether NHE1 contributes to the self-renewal phenotype identified in our screen. To begin to validate NHE1 as a target, we treated Jurkat T-ALL cells with hexamethylene amiloride (HMA), a more potent NHE1 inhibitor (15,17). HMA caused a dose-dependent reduction in colony formation with limited apoptosis (**Supplemental Figure 3**), recapitulating the phenotype observed with amiloride.

We next examined the relevance of NHE1 in human leukemia using the TARGET-ALL-P2 cohort (18), which includes both B-ALL and T-ALL. *SLC9A1* expression was significantly higher in recurrent than primary ALL samples (**Figure 3A**, *p* = 0.0134), and patients with higher *SLC9A1* expression showed poorer overall survival, although this difference did not reach statistical significance (**Figure 3B**; HR = 2.13, 95% CI 1.21-3.75; adjusted *p* = 0.108). NHE1 protein was also detected in T-ALL patient-derived xenografts and human T-ALL cell lines (**Figure 3C**). To directly test the requirement for NHE1, we performed shRNA-mediated knockdown in Jurkat T-ALL cells. Four independent shRNAs targeting *SLC9A1* reduced NHE1 protein expression, with shRNA #1 and shRNA #4 producing the strongest knockdown (**Figure 3D-E**). Reduced *SLC9A1* transcript levels were independently confirmed by RT-qPCR (**Supplemental Figure 4A**). NHE1 knockdown led to a >70% reduction in colony formation compared with scrambled control (**Figure 3F-G**, *p* < 0.0001), whereas the size of colonies that did form was not significantly altered (**Supplemental Figure 4B**). This pattern suggests that NHE1 depletion primarily impairs the ability of T-ALL cells to self-renew and initiate clonogenic growth, rather than simply limiting the expansion of established colonies. Consistent with this effect on self-renewal capacity, shRNA #1 also reduced expression of selected genes previously linked to self-renewal and leukemia cell state (19–23) (**Supplemental Figure 4C**).

**Figure 3.**
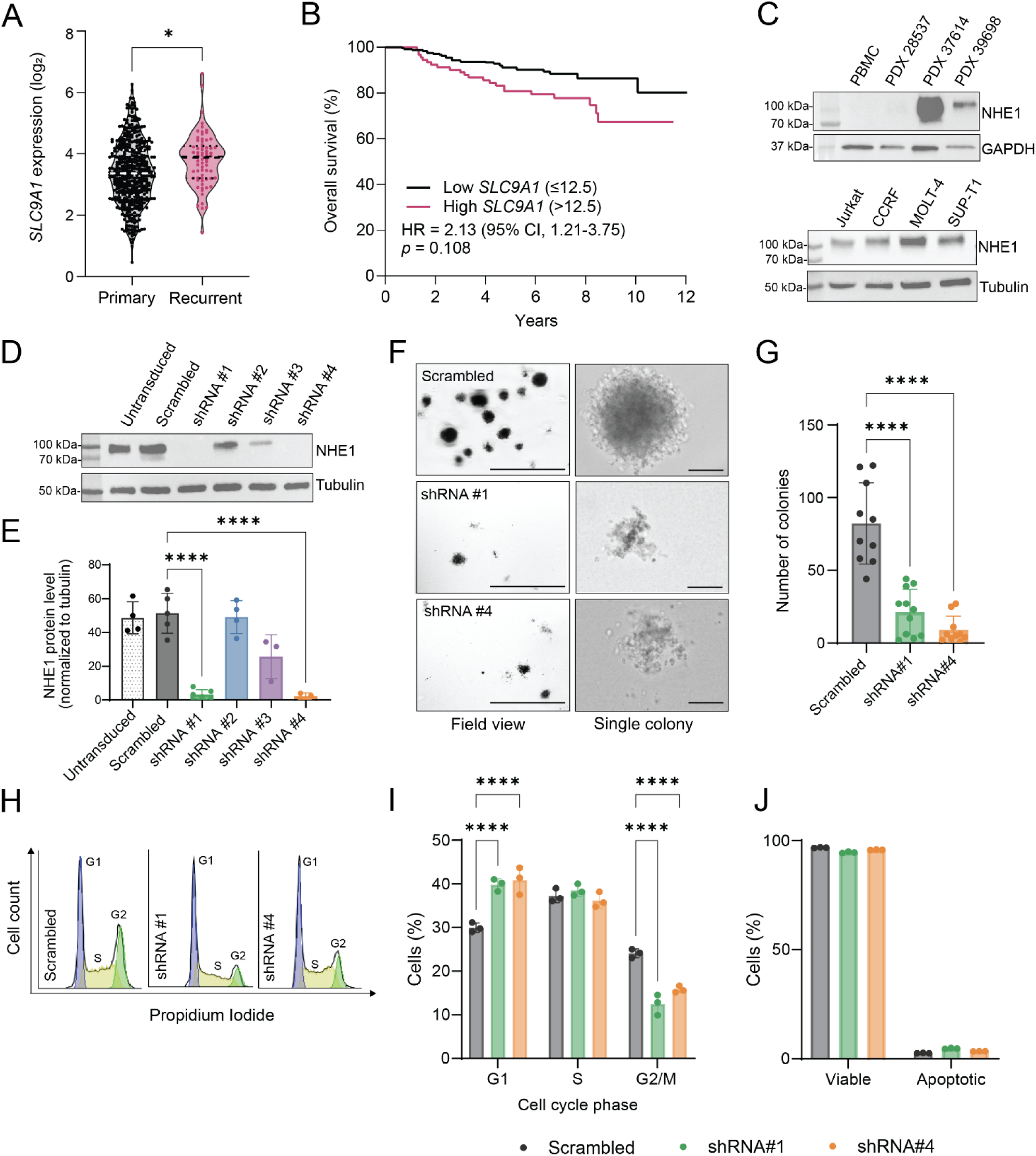
NHE1 depletion suppresses clonogenic growth and induces G1 accumulation in T-ALL cells. **(A)** *SLC9A1* expression in primary (*n* = 463) and recurrent (*n* = 69) ALL samples from the TARGET-ALL-P2 cohort. \*\**p* = 0.0134 by linear mixed-effects model accounting for repeated patient samples. **(B)** Overall survival in TARGET-ALL-P2 patients stratified by *SLC9A1* expression using an optimized cutoff of 12.5 determined by maximally selected rank statistics (maxstat). HR, 2.13 (95% CI, 1.21-3.75); cutoff-adjusted log-rank *p* = 0.108. **(C)** NHE1 protein expression in peripheral blood mononuclear cells (PBMCs), T-ALL patient-derived xenografts, and T-ALL cell lines. GAPDH or tubulin served as loading controls. **(D-E)** NHE1 protein expression in Jurkat cells transduced with scrambled control shRNA or 4 independent shRNAs targeting *SLC9A1*. Representative immunoblot (**D**) and densitometric quantification normalized to tubulin (**E**) are shown. **(F-G)** Colony formation by Jurkat cells expressing scrambled shRNA, shRNA #1, or shRNA #4. Representative field and single-colony images after 14 days (**F**) and colony quantification (**G**) are shown. Scale bars, 1000 μm and 100 μm. **(H-I)** Cell-cycle distribution assessed by propidium iodide staining. Representative DNA-content profiles (**H**) and quantification of cells in G1, S, and G2/M (**I**) are shown. **(J)** Viable and apoptotic cells assessed by Annexin V/propidium iodide staining; apoptotic cells represent combined early– and late-apoptotic populations. Symbols represent independent biological replicates; bars indicate mean ± SD. Statistical significance in **E** and **G** was determined by one-way ANOVA with Dunnett’s multiple-comparisons test and in **I** by two-way ANOVA with Dunnett’s multiple-comparisons test. **** *p* < 0.0001.

NHE1 knockdown also resulted in G_0_/G_1_ accumulation, with corresponding reductions in G_2_/M phases (**Figure 3H-I**, *p* < 0.0001). Despite the pronounced effect on colony formation and cell-cycle distribution, NHE1 depletion caused only a modest increase in apoptosis (**Figure 3J**), indicating that the loss of in vitro self-renewal capacity was not due to cytotoxicity. Notably, the reduction in colony formation, cell cycle changes, and limited apoptosis following genetic depletion of NHE1 closely mirrored the phenotypes observed with amiloride. The concordant pharmacologic and genetic phenotypes identify NHE1 as a previously unrecognized regulator of T-ALL self-renewal and a biologically relevant dependency uncovered by the functional screen.

### NHE1 depletion induces broad transcriptomic and proteomic remodeling in T-ALL cells

To identify cellular programs that might underlie the self-renewal phenotype, we performed bulk RNA sequencing and quantitative proteomic profiling. NHE1 knockdown resulted in widespread changes in gene expression compared with scrambled control (**Figure 4A**). Gene set enrichment analysis (GSEA) demonstrated bidirectional remodeling of oncogenic signaling programs following NHE1 knockdown (**Figure 4B**): KRAS signaling was reduced following NHE1 knockdown (NES = 1.70, FDR = 0.04), whereas MYC Targets V1 showed a trend toward enrichment in NHE1-knockdown cells (NES = −1.59, FDR = 0.06). NOTCH and Hedgehog signaling showed similar trends toward reduction following NHE1 knockdown (**Supplemental Figure 5A**). At the gene level, these transcriptional changes spanned gene involved in cell signaling and cell state, adhesion and migration, and metabolic regulation (**Figure 4C**), with broader changes in extracellular matrix organization, lipid metabolism, transcriptional regulation, and immune/inflammatory responses (**Supplemental Figure 5B**).

**Figure 4.**
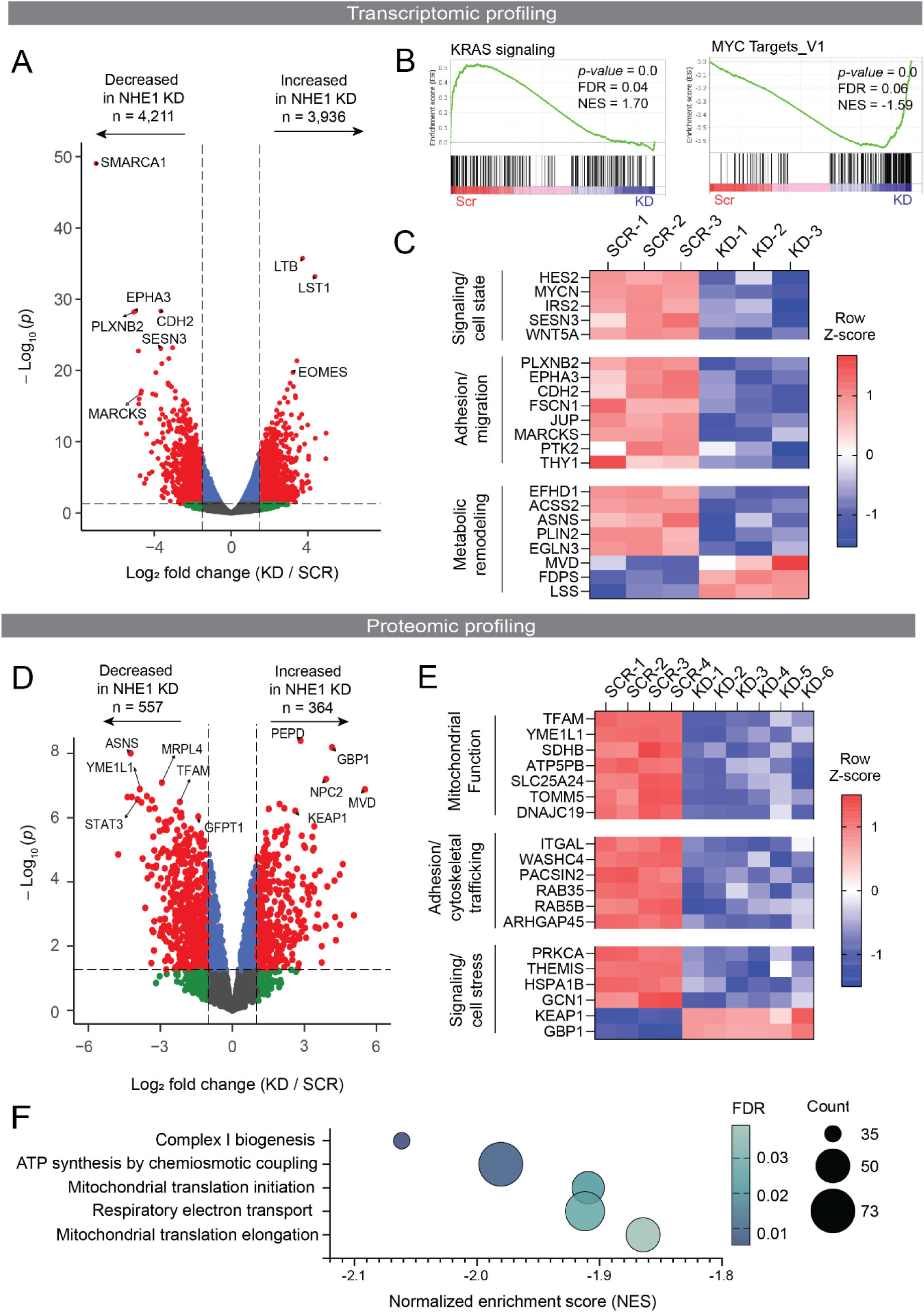
NHE1 depletion induces broad transcriptomic and proteomic remodeling in T-ALL cells. **(A)** Volcano plot of differentially expressed genes in Jurkat cells expressing NHE1 shRNA #1 versus scrambled control. Genes decreased or increased following NHE1 knockdown are shown at left and right, respectively; numbers indicate genes meeting the differential-expression threshold. **(B)** GSEA showing KRAS signaling and MYC Targets V1 gene sets. NES and FDR are indicated; positive NES denotes enrichment in scrambled control and negative NES enrichment in NHE1-knockdown cells. **(C)** Heatmap of selected significantly altered genes associated with signaling/cell state, adhesion/migration, and metabolic remodeling. Expression is shown as row Z-scores (*n* = 3/group). **(D)** Volcano plot of quantitative proteomic changes following NHE1 knockdown. Proteins decreased or increased in NHE1-knockdown cells are shown at left and right, respectively; numbers indicate proteins meeting the differential-expression threshold. **(E)** Heatmap of selected significantly altered proteins associated with mitochondrial function, adhesion/cytoskeletal trafficking, and signaling/stress response. VSN-normalized protein abundance is displayed as row Z-scores (scrambled, *n* = 4; NHE1 knockdown, *n* = 6). **(F)** Reactome pathways significantly altered following NHE1 knockdown (FDR < 0.05). Bubble position indicates normalized enrichment score (NES), size indicates the number of proteins represented in each pathway, and color indicates FDR. Negative NES values indicate pathways enriched for proteins decreased following NHE1 knockdown.

Complementing the RNA sequencing data, proteomic profiling identified broad changes in protein abundance following NHE1 knockdown (**Figure 4D**), including decreased proteins involved in mitochondrial function and homeostasis, as well as changes in proteins involved in cell adhesion and signaling/stress responses (**Figure 4E**). Pathway enrichment analysis revealed a particularly strong mitochondrial signature. All Reactome pathways meeting FDR < 0.05 had negative enrichment scores, indicating coordinated depletion of proteins involved in mitochondrial respiration, ATP production, and mitochondrial protein synthesis (**Figure 4F; Supplemental Figure 6**). Independent KEGG, WikiPathways, and Gene Ontology analyses converged on the same pattern, identifying reduced oxidative phosphorylation and mitochondrial energy-production programs (**Supplemental Figure 6**). The convergence of the transcriptomic and proteomic datasets on mitochondrial pathways identified mitochondrial fitness as a candidate cellular process that could contribute to the NHE1-dependent self-renewal phenotype.

### NHE1 inhibition and depletion impair mitochondrial fitness in T-ALL cells

To functionally validate the mitochondrial defects identified by transcriptomic and proteomic profiling, we performed Seahorse mitochondrial stress testing following pharmacologic or genetic NHE1 perturbation. This assay measures oxygen consumption in living cells and quantifies basal respiration, ATP-linked respiration, and spare respiratory capacity, reflecting the ability of cells to meet basal and increased energetic demands. Amiloride significantly reduced all three parameters in Jurkat cells (**Figure 5A**; all *p* < 0.001), with similar effects in CCRF-CEM and MOLT-4 cells (**Supplemental Figure 7A-B**). A second NHE1 inhibitor, HMA, also significantly reduced basal and ATP-linked respiration in Jurkat cells (**Supplemental Figure 7C**). NHE1 depletion with either shRNA #1 or shRNA #4 similarly reduced basal respiration, ATP-linked respiration, and spare respiratory capacity (**Figure 5B**; all *p* < 0.0001). These findings functionally validated the mitochondrial bioenergetic defects associated with NHE1 loss that were predicted by the omics analyses.

**Figure 5.**
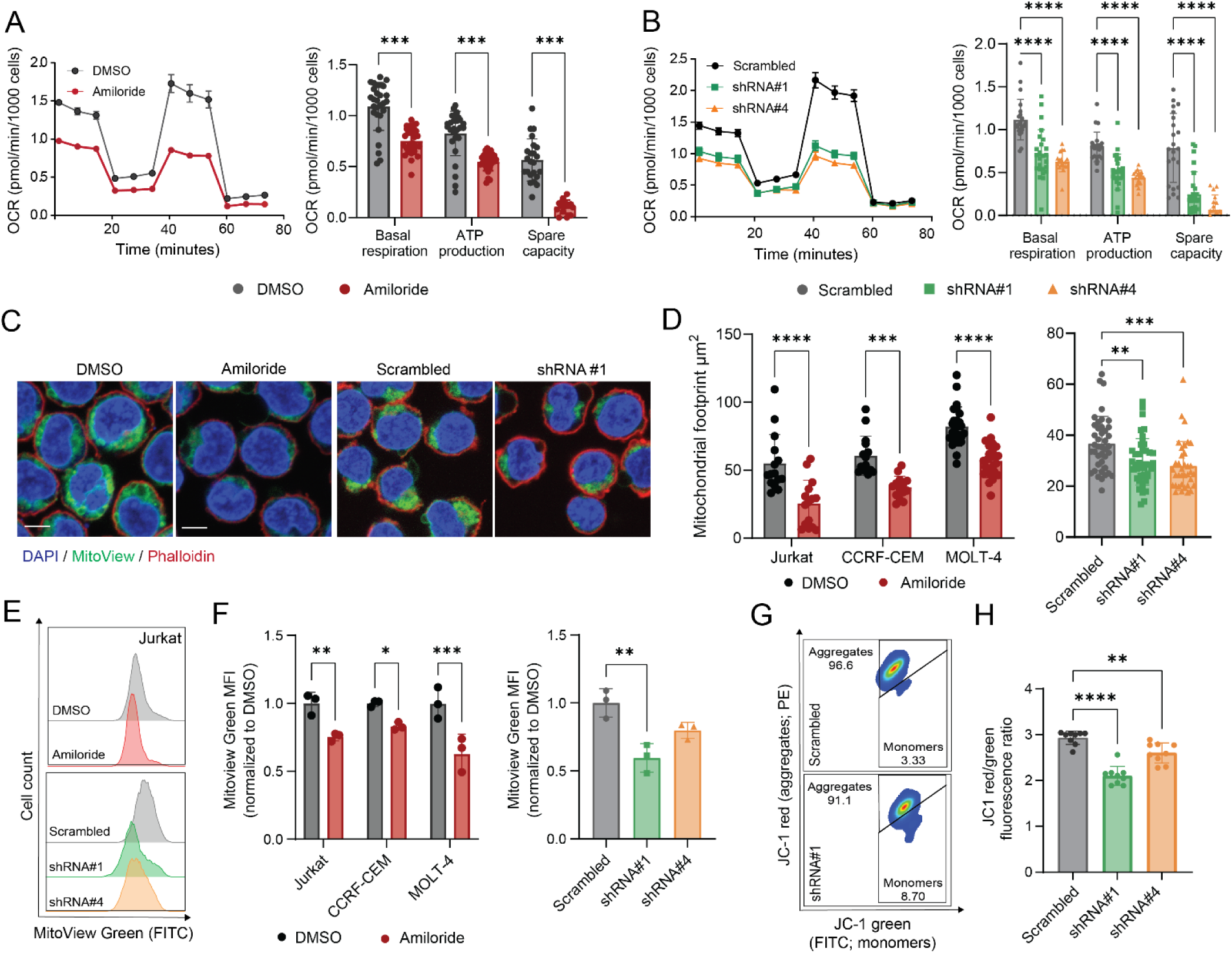
Pharmacologic inhibition and genetic depletion of NHE1 impair mitochondrial function and content in T-ALL cells. **(A)** Mitochondrial stress-test oxygen consumption rate (OCR) profiles and quantification of basal respiration, ATP production, and spare respiratory capacity in Jurkat cells treated with DMSO or amiloride (100 μM, 24 hours). **(B)** Mitochondrial stress-test OCR profiles and quantification of basal respiration, ATP production, and spare respiratory capacity in Jurkat cells expressing scrambled control, NHE1 shRNA #1, or NHE1 shRNA #4. **(C)** Representative confocal images of CCRF-CEM cells treated with DMSO or amiloride (100 μM, 24 hours) and Jurkat cells expressing scrambled control or NHE1 shRNA #1, stained with DAPI (blue), MitoView Green (green), and phalloidin (red). Scale bar, 10 μm. **(D)** Quantification of mitochondrial footprint in Jurkat, CCRF-CEM, and MOLT-4 cells treated with DMSO or amiloride (left) and in Jurkat cells expressing scrambled control, NHE1 shRNA #1, or NHE1 shRNA #4 (right). Mitochondrial footprint was quantified using the MiNA (Mitochondrial Network Analysis) ImageJ plugin. **(E)** Representative flow-cytometry histograms of MitoView Green fluorescence in Jurkat cells treated with DMSO or amiloride (top) and in Jurkat cells expressing scrambled control, NHE1 shRNA #1, or NHE1 shRNA #4 (bottom). **(F)** Quantification of MitoView Green mean fluorescence intensity (MFI), normalized to the corresponding control, in Jurkat, CCRF-CEM, and MOLT-4 cells treated with DMSO or amiloride (left) and in Jurkat cells expressing scrambled control, NHE1 shRNA #1, or NHE1 shRNA #4 (right). **(G)** Representative JC-1 flow-cytometry plots showing red JC-1 aggregates (PE) and green JC-1 monomers (FITC) in Jurkat cells expressing scrambled control or NHE1 shRNA #1. **(H)** Quantification of the JC-1 red/green fluorescence ratio in Jurkat cells expressing scrambled control, NHE1 shRNA #1, or NHE1 shRNA #4 as a measure of mitochondrial membrane potential. Data are shown as mean ± SD with individual data points overlaid. Pharmacologic Seahorse comparisons were assessed by mixed-effects analysis with Sidak’s multiple-comparisons test (**A**), and shRNA Seahorse comparisons by mixed-effects analysis with Dunnett’s multiple-comparisons test (**B**). Mitochondrial-footprint comparisons were assessed by two-way ANOVA with Sidak’s multiple-comparisons test for drug treatment and one-way ANOVA with Dunnett’s multiple-comparisons test for shRNA conditions (**D**). MitoView Green pharmacologic comparisons were assessed by two-way repeated-measures ANOVA with Sidak’s multiple-comparisons test, and shRNA comparisons by one-way repeated-measures ANOVA with Dunnett’s multiple-comparisons test (**F**). * *p* < 0.05, ** *p* < 0.01, *** *p* < 0.001, **** *p* < 0.0001.

Because proteomic profiling also identified reduced abundance of proteins involved in mitochondrial homeostasis, we next examined mitochondrial content. Confocal imaging of MitoView Green-labeled mitochondria was used to quantify mitochondrial footprint, defined as the cellular area occupied by mitochondrial signal. Mitochondrial footprint was significantly reduced following amiloride treatment in Jurkat, CCRF-CEM, and MOLT-4 cells and following NHE1 knockdown in Jurkat cells (**Figure 5C-D**). Flow-cytometric analysis of MitoView Green across larger cell populations provided an independent measure of mitochondrial abundance and likewise demonstrated reduced mitochondrial content following amiloride treatment across all three T-ALL cell lines and in NHE1 knockdown cells (**Figure 5E-F**).

Finally, we examined mitochondrial membrane potential (Δψm), the electrochemical gradient across the inner mitochondrial membrane required for efficient ATP synthesis. JC-1 staining demonstrated a reduced red-to-green fluorescence ratio in both NHE1-knockdown lines compared with scrambled control (**Figure 5G-H**; shRNA #1, *p* < 0.0001; shRNA #4, *p* < 0.01), consistent with a significantly decreased mitochondrial membrane potential. Together, these data indicate that diminished NHE1 activity impairs mitochondrial fitness in T-ALL cells, with lower respiratory capacity, mitochondrial content, and membrane potential.

### NHE1 depletion alters the metabolic state of T-ALL cells

To determine whether the mitochondrial defects associated with NHE1 depletion were accompanied by broader metabolic consequences, we performed GC-MS metabolomic profiling comparing NHE1 shRNA#1 cells with scrambled controls. Principal component analysis (PCA) demonstrated clear separation of NHE1-knockdown and control samples, with PC1 accounting for 59.3% of the variance, indicating broad remodeling of the metabolic state following NHE1 depletion (**Figure 6A**). Consistent with this separation, differential abundance analysis identified a subset of metabolites with marked changes following NHE1 depletion (**Figure 6B**). We next focused on metabolic changes that were particularly relevant to the mitochondrial phenotype identified in **Figure 5**. The tricarboxylic acid (TCA) cycle intermediates citrate and α-ketoglutarate were significantly reduced in NHE1-knockdown cells (*q* = 0.0117 and *q* = 0.0151, respectively), as was pantothenic acid, a precursor required for coenzyme A metabolism (*q* = 0.0218) (**Figure 6C**). In contrast, L-aspartate, which links mitochondrial and cytosolic metabolism, and myo-inositol phosphate, a component of inositol phosphate metabolism, were significantly increased (*q* = 0.0155 and *q* = 0.0340, respectively), demonstrating that NHE1 depletion altered multiple interconnected metabolic programs. Visualization of all 68 identified metabolites further demonstrated differences spanning amino acid, carbohydrate, lipid, nucleotide, polyamine, and TCA cycle-associated metabolism (**Figure 6D**). These metabolomic changes extend the mitochondrial phenotype identified in Figure 5, showing that NHE1 loss produces broader metabolic remodeling that could contribute to impaired T-ALL self-renewal.

**Figure 6.**
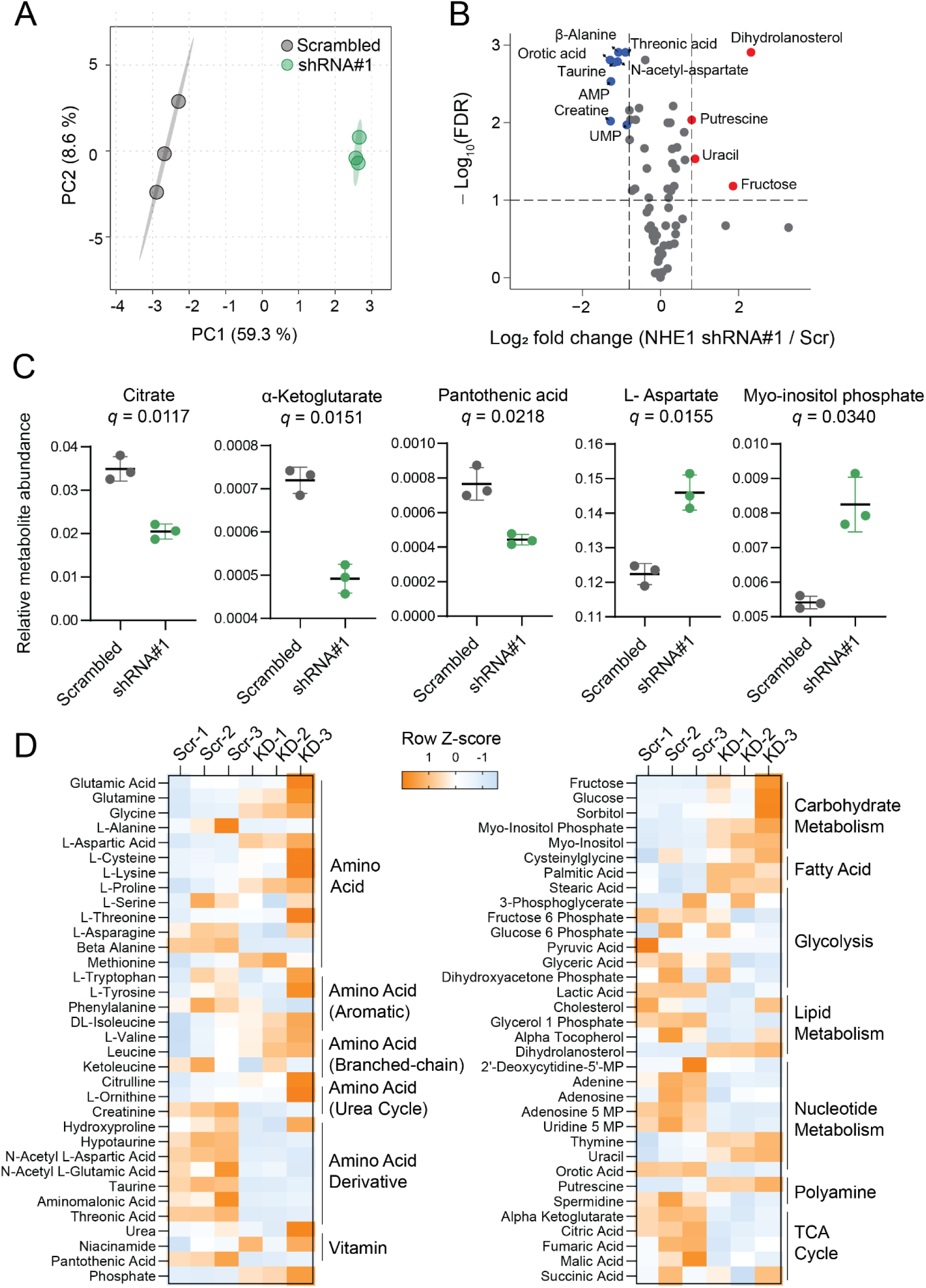
NHE1 depletion alters the metabolic state of human T-ALL cells. **(A)** Principal component analysis (PCA) of GC-MS metabolomic profiles from Scrambled control and NHE1 shRNA#1 cells. Each point represents an independent biological replicate (*n* = 3 per group). **(B)** Volcano plot showing differential metabolite abundance in NHE1 shRNA#1 cells relative to Scrambled control. Metabolites meeting log₂ fold change > 0.8 or <-0.8 and FDR < 0.1 are highlighted; blue indicates decreased abundance and red indicates increased abundance in NHE1-depleted cells. Selected significant metabolites are labeled. **(C)** Relative abundance of representative metabolites significantly altered following NHE1 depletion. Points represent independent biological replicates; horizontal lines indicate mean ± SD. *q* values were determined using multiple unpaired comparisons of log-transformed metabolite abundance with Benjamini-Hochberg false discovery rate correction. **(D)** Heatmap showing row Z-scores for identified metabolites across Scrambled control and NHE1 shRNA#1 samples, grouped by metabolic class. Orange indicates higher and blue indicates lower relative abundance for each metabolite.

### NHE1 depletion reduces T-ALL burden in hematopoietic tissues in vivo

Finally, we asked whether the cellular fitness associated with NHE1 depletion was reflected in leukemia behavior in vivo. Luciferase-expressing Jurkat cells expressing scrambled control or NHE1-targeting shRNA#1 were transplanted into immunodeficient mice. Whole-body bioluminescence increased over time in both groups with similar overall trajectories (**Figure 7A-B**), and survival was not significantly different between mice receiving scrambled control and shRNA#1 cells (**Figure 7C**; *p* = 0.47). Thus, NHE1 knockdown did not substantially alter overall leukemia progression in this model.

**Figure 7.**
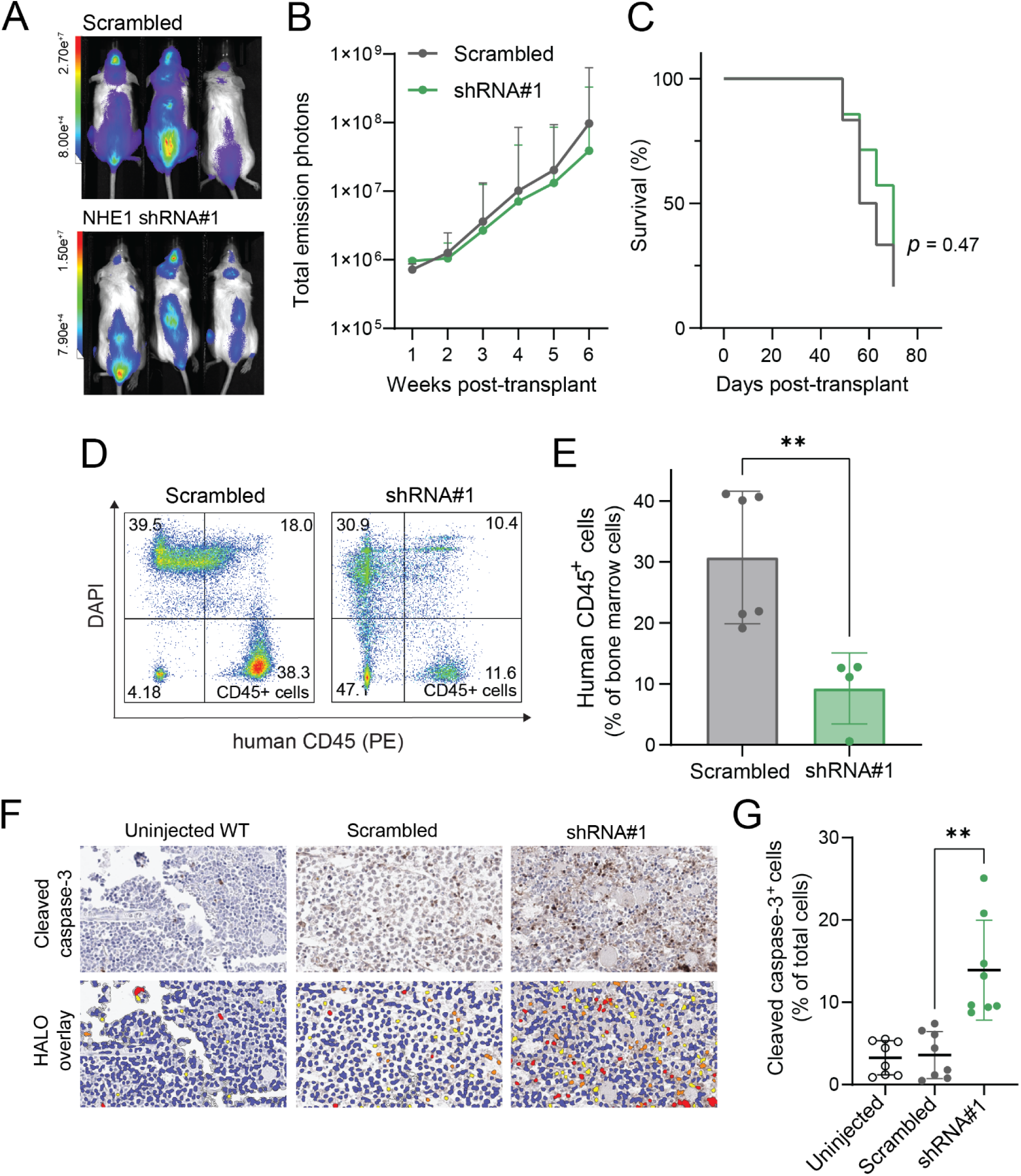
NHE1 knockdown reduces T-ALL burden in the bone marrow in vivo. **(A)** Representative whole-body bioluminescence images of immunodeficient mice transplanted by tail-vein injection with luciferase-expressing Jurkat cells expressing scrambled control or NHE1-targeting shRNA#1 at 6 weeks post-transplantation. **(B)** Longitudinal bioluminescence imaging of mice transplanted with scrambled control or shRNA#1 cells over 6 weeks. Total emission photons are displayed on a logarithmic scale. Data are shown as geometric mean with geometric SD. **(C)** Kaplan-Meier survival analysis of mice transplanted with scrambled control or shRNA#1 cells. Statistical significance was assessed by log-rank test (*p* = 0.47). **(D)** Representative flow-cytometry plots of bone marrow cells stained with DAPI and PE-conjugated human CD45. Viable human leukemia cells were identified as DAPI⁻/human CD45⁺ cells. **(E)** Quantification of human CD45⁺ cells in the bone marrow of mice transplanted with scrambled control or shRNA#1 cells. Each point represents one mouse; data are shown as mean ± SD. Statistical significance was assessed using a two-tailed Mann–Whitney test. **(F)** Representative cleaved caspase-3 immunohistochemistry of bone marrow sections from uninjected wild-type (WT) mice and mice transplanted with scrambled control or shRNA#1 cells. Corresponding HALO nuclear phenotyper overlays are shown below; yellow, orange, and red indicate weak, moderate, and strong cleaved caspase-3 staining, respectively. Scale bars, 50 μm. **(G)** HALO-based quantification of cleaved caspase-3-positive cells as a percentage of total cells in bone marrow sections from uninjected WT mice and mice transplanted with scrambled control or shRNA#1 cells. data are shown as mean ± SD. Statistical significance was assessed using a Kruskal–Wallis test with Dunn’s multiple-comparisons test. \**p* < 0.01

Despite similar whole-body leukemia burden, mice transplanted with NHE1-depleted cells had significantly fewer viable human CD45-positive cells in the bone marrow than scrambled controls (**Figure 7D-E**). Because the bone marrow is a major site of T-ALL localization and persistence (24,25), this finding suggests that NHE1 loss compromises leukemia fitness within the marrow rather than simply slowing leukemia growth. Bone marrow from mice receiving shRNA#1 cells also contained a significantly greater proportion of cleaved caspase-3-positive cells than marrow from scrambled controls (**Figure 7F-G**), consistent with increased apoptotic activity in the marrow environment.

A similar reduction in leukemia-associated tissue involvement was observed in the spleen. Mice receiving scrambled control cells developed enlarged, heavier spleens with prominent leukemic blasts and marked disruption of normal splenic tissue, whereas spleens from mice receiving shRNA#1 cells were smaller and more closely resembled those from uninjected mice (**Supplemental Figure 8A-D**). Consistent with this, eosin-positive tissue area was markedly reduced in scrambled-control spleens but preserved in the shRNA#1 group at levels similar to uninjected mice, consistent with better preservation of normal spleen tissue (**Supplemental Figure 8E**). These findings show that NHE1 loss has relatively little effect on overall leukemia progression but substantially reduces T-ALL involvement in hematopoietic tissues, extending the self-renewal and metabolic phenotypes identified in vitro to a distinct loss of leukemia fitness in vivo.

## Discussion

Our findings identify the sodium/hydrogen exchanger NHE1 as a previously unrecognized regulator of leukemia-propagating activity in T-ALL. Because T-ALL LPCs lack reliable defining surface markers (3,4), we used an in vivo screening strategy that measured their ability to regenerate leukemia following transplantation, without requiring their prior identification or isolation. Zebrafish T-ALL clones with an unusually high LPC frequency (∼10%) allowed us to assess leukemia propagation using as few as 25 cells per recipient, while the scalability of zebrafish transplantation enabled us to evaluate 786 FDA-approved compounds across more than 3,000 animals. Unlike conventional drug screens that primarily measure bulk cell proliferation or viability, this approach identified compounds that impaired leukemia-propagating capacity in vivo. Among these, amiloride markedly reduced LPC frequency without substantially delaying leukemia development at higher transplantation doses. Genetic depletion of NHE1 also impaired in vitro self-renewal capacity, with only modest effects on cell growth and apoptosis. These findings highlight how leukemia-propagating function can be impaired without corresponding effects on bulk leukemia growth or survival, a distinction that may be missed by conventional screening approaches.

Previous studies have implicated NHE1 in leukemia proliferation, differentiation, and therapeutic resistance across several hematologic malignancies, including AML, CML, and T-ALL(11–15). In AML, Hyun et al. reported higher NHE1 expression in CD34⁺CD38⁻ leukemia stem-cell-enriched populations compared with CD34⁺CD38⁺ cells. Pharmacologic inhibition of NHE1 also induced G1 cell-cycle arrest and, at higher concentrations, apoptosis(12). Although these observations suggested a potential role for NHE1 in leukemia stem-cell biology, they did not establish whether NHE1 was required for leukemia-propagating activity. Our study addresses this gap by combining genetic evidence of impaired in vitro self-renewal capacity with direct measurements of leukemia-propagating frequency in vivo. These findings support a role for NHE1 to T-ALL propagation and provides a rationale for investigating the cellular mechanisms underlying this dependency.

Mitochondrial dysfunction was among the most prominent consequences of NHE1 depletion, with reduced impaired mitochondrial respiration, reduced mitochondrial content and membrane potential, and broad changes in cellular metabolism. Alterations in TCA cycle intermediates, coenzyme A precursor metabolism, and amino acid metabolism suggest that NHE1 loss affects multiple pathways involved in mitochondrial energy production. The potential significance of these changes for leukemia propagation is supported by studies in AML and CML. In AML, Jones et al. showed that leukemia stem cells depend on amino acid metabolism to support oxidative phosphorylation (26), while Kuntz et al. demonstrated that inhibiting mitochondrial protein translation, in combination with imatinib, could eliminate therapy-resistant CML stem cells(27). This is particularly relevant to our finding that NHE1 depletion reduces the abundance of proteins involved in mitochondrial translation. Mitochondrial metabolism is also important in T-ALL, where Baran et al. demonstrated that inhibition of mitochondrial complex I suppresses NOTCH1-driven leukemia growth (28). Given these established metabolic dependencies, the mitochondrial defects associated with NHE1 loss may contribute to its effects on leukemia-propagating activity.

NHE1 regulates intracellular pH through the exchange of intracellular H⁺ for extracellular Na⁺, a process that can influence metabolic enzyme activity, substrate utilization, and cellular signaling (15). NHE1 can also affect calcium homeostasis through functional interactions with the Na⁺/Ca²⁺ exchanger NCX1(29). Calcium-dependent processes have previously been implicated in leukemia-propagating activity. For example, In AML, Sheth et al. demonstrated that mitochondrial calcium uptake supports oxidative phosphorylation and leukemia stem-cell function, with inhibition of calcium uptake impairing mitochondrial metabolism and leukemia-propagating activity(30). In T-ALL, Gachet et al. showed that calcineurin, a calcium-dependent phosphatase, is required for leukemia propagation (31). These studies suggest that ion homeostasis may influence both mitochondrial fitness and leukemia-propagating activity, providing a potential biological explanation for the effects of NHE1 depletion in T-ALL.

NHE1 depletion also reduced leukemia involvement in the bone marrow and spleen despite similar overall leukemia progression. Increased apoptotic activity in the marrow suggests that NHE1 may contribute to leukemia-cell survival within this environment. The bone marrow contains distinct microenvironments with varying extracellular pH, and recent studies in AML have demonstrated that these pH differences influence leukemia-initiating cell localization and activity (32). As a major regulator of intracellular pH, NHE1 may help T-ALL cells adapt to the varying conditions encountered within the marrow. In addition, our transcriptomic and proteomic analyses identified changes in pathways associated with cell adhesion, migration, and signaling, suggesting that NHE1 may also influence leukemia interactions with the microenvironment. NHE1 has established roles in cell polarization and directed migration through its interactions with the actin cytoskeleton(33–35), and previous studies in T-ALL demonstrated that NHE1 inhibition or silencing impairs CCL25-induced migration(14). These observations raise the possibility that NHE1 supports leukemia-propagating activity not only through its effects on cellular metabolism, but also by facilitating leukemia-cell interactions with supportive hematopoietic niches.

The limitations of this study include the lack of direct assessment of leukemia-propagating cell frequency following genetic NHE1 depletion in vivo and uncertainty regarding whether mitochondrial dysfunction or altered interactions with the bone marrow microenvironment directly contribute to leukemia-propagating cell activity. Further studies in patient-derived T-ALL models will also be important for establishing the therapeutic relevance of NHE1 inhibition. Nevertheless, our findings identify NHE1 as a previously unrecognized contributor to T-ALL leukemia-propagating function and reveal its importance for mitochondrial fitness and leukemia involvement in hematopoietic tissues. These observations suggest a broader role for ion transport in sustaining the cellular functions that allow T-ALL to propagate and persist, opening a new direction for investigating vulnerabilities in leukemia-propagating cells.

## Acknowledgements

Funding for this research was provided by the National Cancer Institute (R37CA227656 to J.S.B.) and the Kentucky Pediatric Cancer Research Trust Fund (PON2 728 2400001524 to J.S.B.). This research was also supported by the Redox Metabolism, Flow Cytometry and Immune Monitoring, and Biostatistics and Bioinformatics Shared Resources of the University of Kentucky Markey Cancer Center (P30CA177558) The authors thank Eva Aldarondo for zebrafish husbandry and maintenance, Emma Arvin for general research assistance, Dr. Benjamin Burke for helpful discussions regarding proteomics analysis, and Dr. Viral Oza for assistance with transcriptomics analysis.

## Authorship Contributions

Conceptualization, M.A.A. and J.S.B.; data curation, M.A.A. and D.H.; formal analysis, M.A.A., D.H., C.W., R.C.B. and J.S.B; funding acquisition, J.S.B.; investigation, M.A.A., M.G.H., Y.C., T.C.C., M.G.P., J.T.J., D.H., L.T.T., and R.C.B.; methodology, M.A.A., M.G.H., Y.C., and R.C.B.; project administration, M.A.A. and J.S.B.; resources, C.W., R.C.B., and J.S.B.; supervision, C.W. and J.S.B.; visualization, M.A.A. and J.S.B.; writing – original draft, M.A.A.; writing – review & editing, J.S.B. All authors reviewed and approved the final version of the manuscript.

## Scientific Category

Lymphoid Neoplasia

## Data Sharing Statement

RNA sequencing data are available through the Gene Expression Omnibus under accession number **GSE341718**. For other original data, please contact Jessica S. Blackburn at.

