## Supplementary material for "NHE1 sustains leukemia-propagating capacity and mitochondrial fitness in T-cell acute lymphoblastic leukemia": Sup Figures 1-8 and Sup Tables 2-4

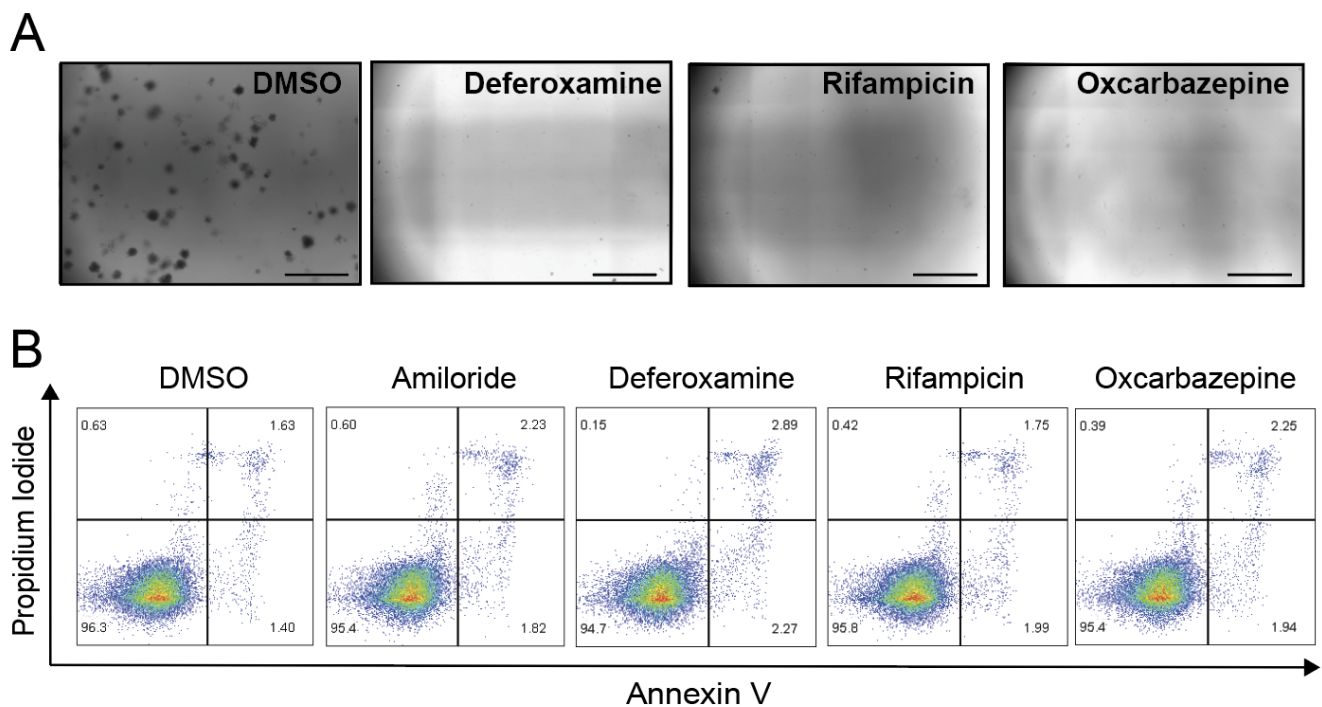

**Supplemental Figure 1. Prioritized compounds suppress T-ALL colony formation while preserving cell viability.** (A) Representative images of Jurkat colonies after 14 days of growth in MethoCult with DMSO or the indicated compounds (10 μM). Scale bar, 2 mm. (B) Representative flow cytometry plots of Annexin V-FITC/propidium iodide (PI) staining in Jurkat cells treated with DMSO or the indicated compounds (10 μM) for 24 h.

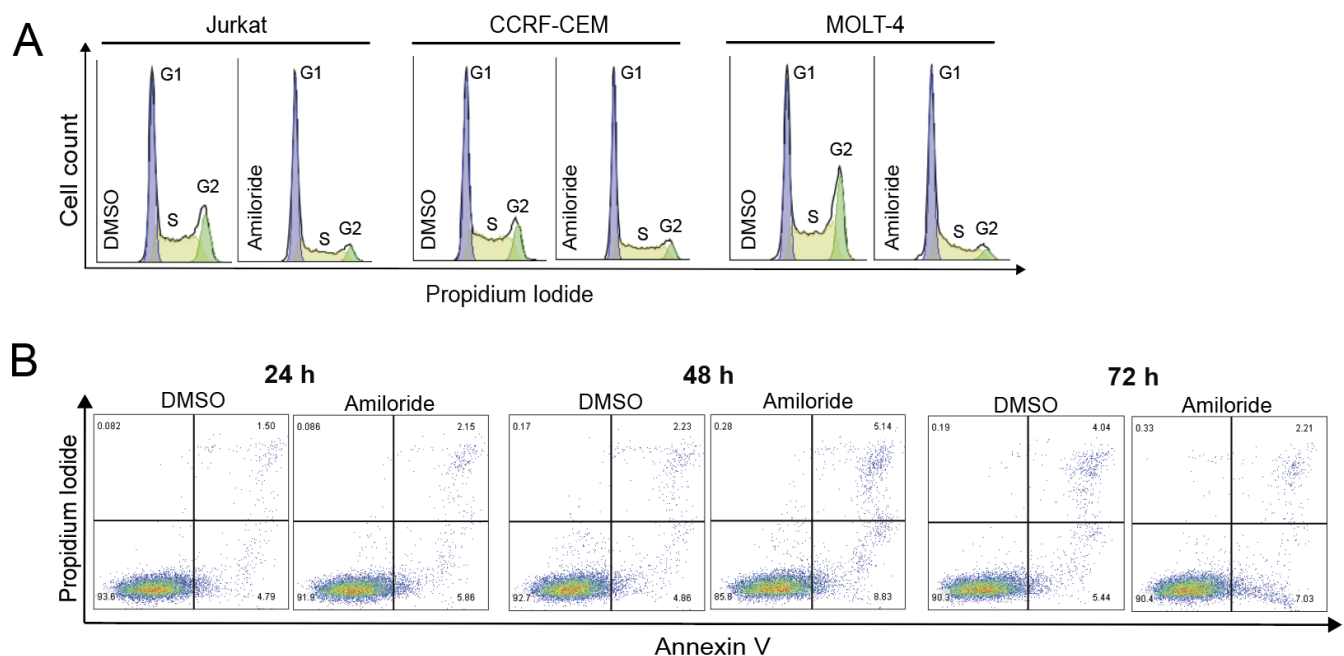

**Supplemental Figure 2. Representative flow cytometry profiles for cell-cycle and apoptosis analyses. (A)** Representative propidium iodide cell-cycle histograms of Jurkat, CCRF-CEM, and MOLT-4 cells following 24-hour treatment with DMSO or 100 μM amiloride. **(B)** Representative Annexin V/propidium iodide flow cytometry plots of Jurkat cells following treatment with DMSO or 100 μM amiloride for 24, 48, or 72 hours.

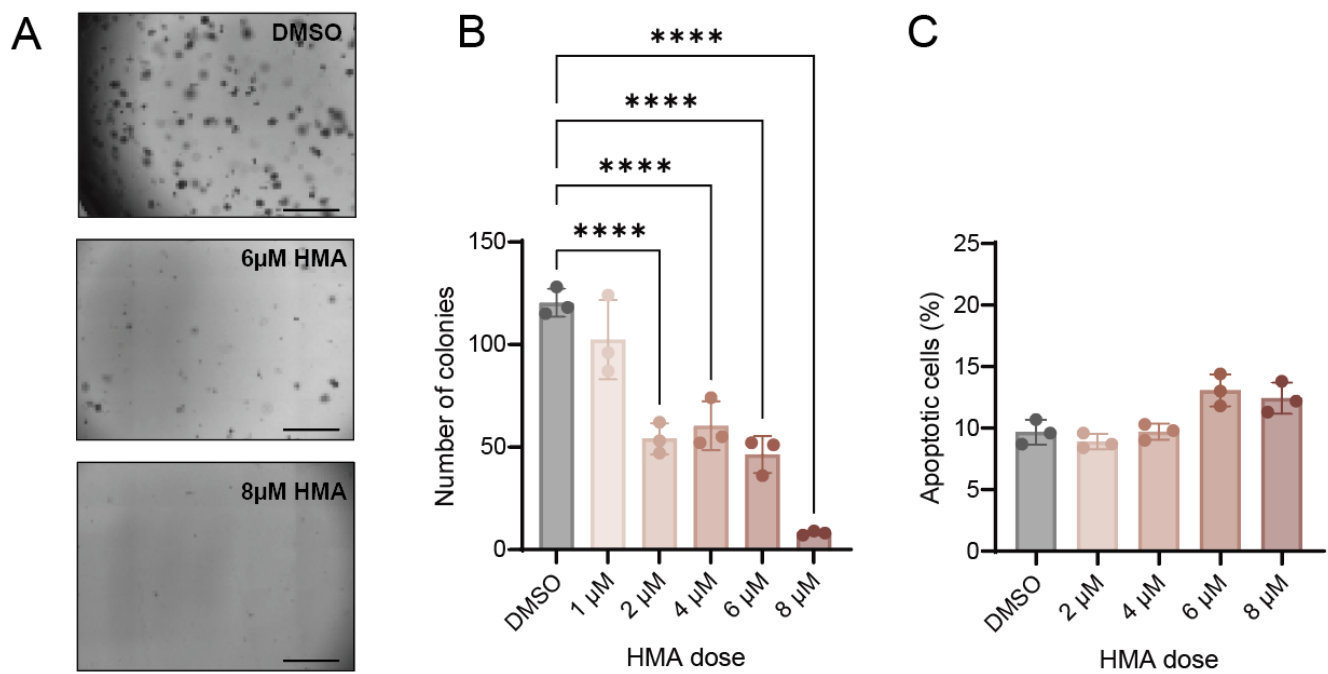

**Supplemental Figure 3. Pharmacologic NHE1 inhibition suppresses T-ALL clonogenic growth with limited acute apoptosis.** (A) Representative field images of Jurkat T-ALL colonies following treatment with DMSO or hexamethylene amiloride (HMA; 6 or 8 μM) after 14 days of colony growth. Scale bars, 2000 μm. (B) Colony formation following treatment with DMSO or increasing concentrations of HMA (1–8 μM). (C) Apoptosis following 24-hour treatment with DMSO or HMA (2–8 μM), assessed by Annexin V-FITC/PI staining. Apoptotic cells represent the combined early- and late-apoptotic populations. Symbols represent independent biological replicates; bars indicate mean ± SD. Statistical significance was determined by one-way ANOVA with Dunnett's multiple-comparisons test versus DMSO. \*\*\*\* $P < 0.0001$.

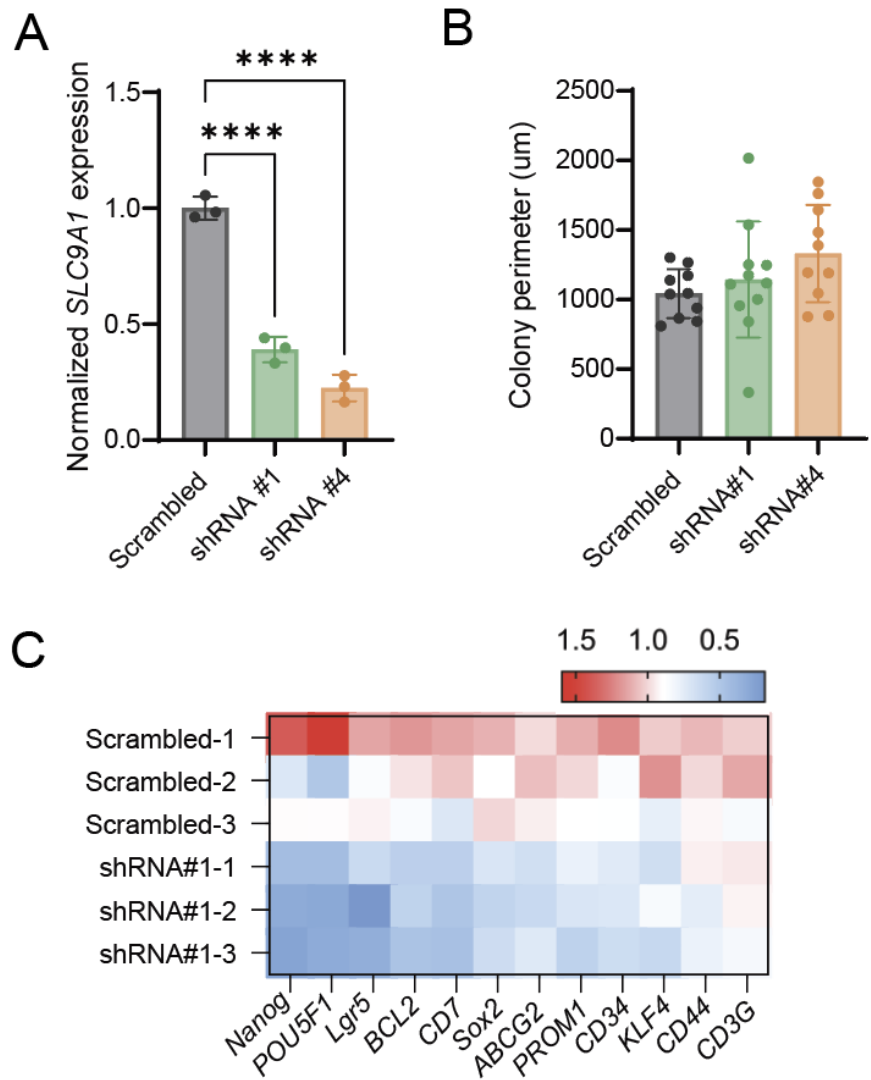

**Supplemental Figure 4. Molecular characterization of NHE1-depleted T-ALL cells.** (A) RT-qPCR analysis of *SLC9A1* expression in Jurkat cells expressing scrambled control shRNA or independent *SLC9A1*-targeting shRNAs #1 and #4. (B) Colony perimeter following colony-forming assays in scrambled control, shRNA#1, and shRNA #4 cells. (C) RT-qPCR analysis of selected genes implicated in self-renewal and leukemia cell state in scrambled control and shRNA #1 cells. Relative transcript abundance was determined by the $\Delta\Delta C_t$ method following normalization to reference genes and is expressed relative to scrambled control. Symbols represent independent biological replicates; bars indicate mean ± SD. Statistical significance was determined by one-way ANOVA with Dunnett's multiple-comparisons test. \*\*\*\* $p < 0.0001$.

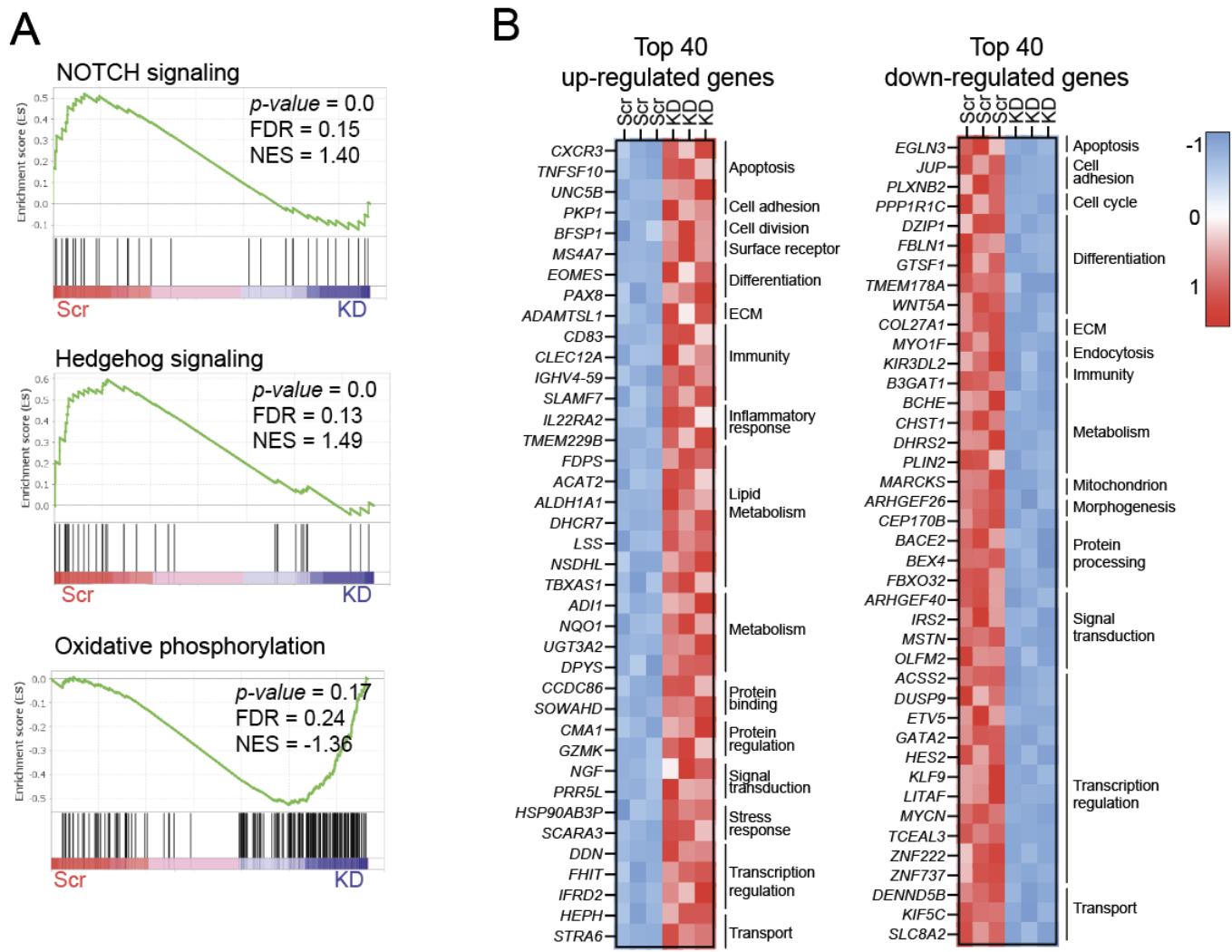

**Supplemental Figure 5. NHE1 depletion alters transcriptional programs in T-ALL cells.** (A) GSEA of RNA-seq data from NHE1-knockdown and scrambled-control Jurkat cells showing NOTCH signaling, Hedgehog signaling, and oxidative phosphorylation gene sets. NES and FDR are indicated; positive NES denotes enrichment in scrambled control and negative NES enrichment in NHE1-knockdown cells. (B) Heatmaps of the top 40 genes increased or decreased following NHE1 knockdown, grouped by functional annotation using UniProtKB and Gene Ontology biological processes. Expression is shown as row Z-scores (n = 3/group).

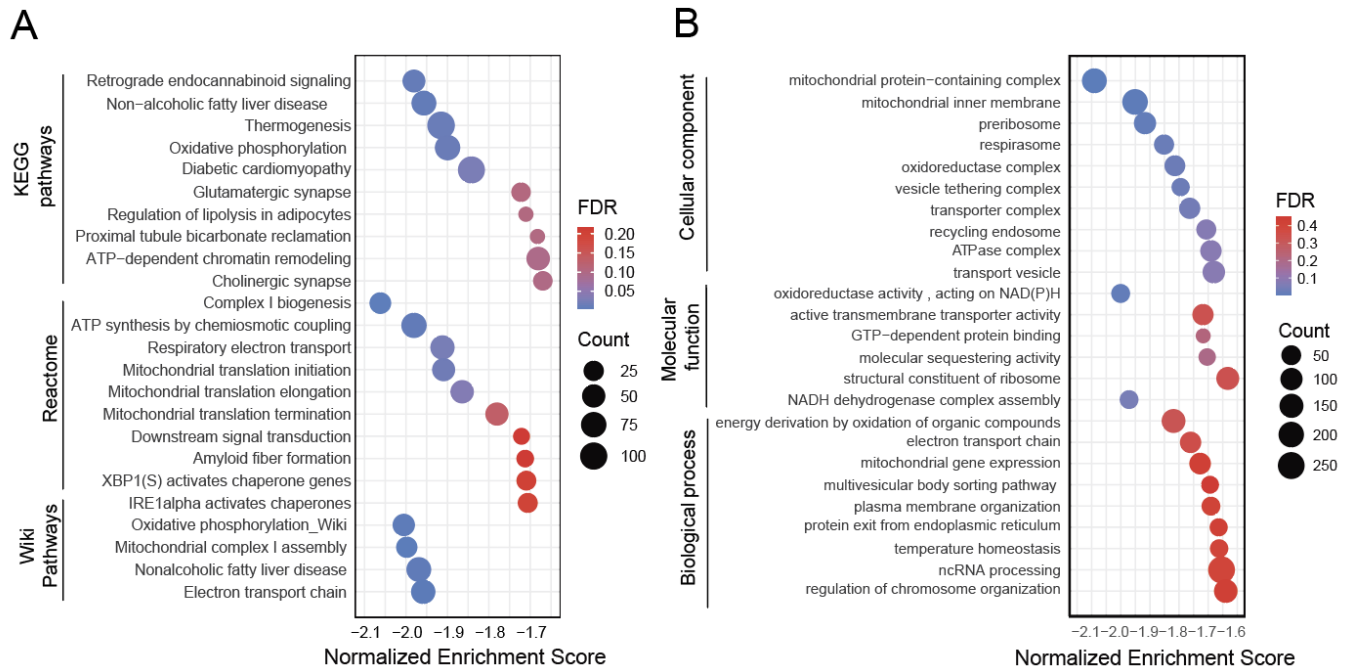

**Supplemental Figure 6. Proteomic pathway analysis identifies coordinated depletion of mitochondrial programs following NHE1 knockdown. (A)** KEGG, Reactome, and WikiPathways enrichment analysis of quantitative proteomic data from NHE1-knockdown versus scrambled-control Jurkat cells. **(B)** Gene Ontology enrichment showing selected cellular component, molecular function, and biological process terms. For both panels, bubble position indicates normalized enrichment score (NES), bubble size indicates protein count, and color indicates FDR.

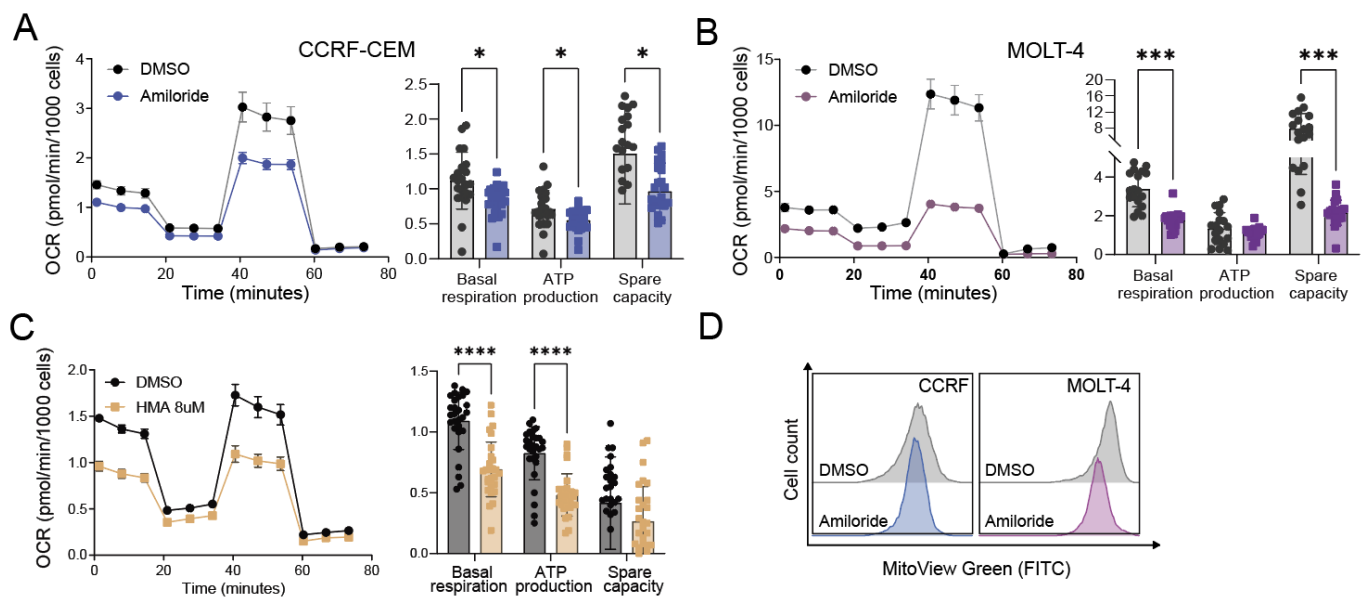

**Supplemental Figure 7. Pharmacologic inhibition of NHE1 reduces mitochondrial respiration and mitochondrial content in T-ALL cells. (A-B)** Mitochondrial stress-test oxygen consumption rate (OCR) profiles and quantification of basal respiration, ATP production, and spare respiratory capacity in CCRF-CEM (**A**) and MOLT-4 (**B**) cells treated with DMSO or amiloride (100 μM, 24 hours). (**C**) Mitochondrial stress-test OCR profile and quantification of basal respiration, ATP production, and spare respiratory capacity in Jurkat cells treated with DMSO or hexamethylene amiloride (HMA; 8 μM, 24 hours). (**D**) Representative flow cytometry histograms of MitoView Green fluorescence in CCRF-CEM and MOLT-4 cells treated with DMSO or amiloride (100 μM, 24 hours). Data are shown as mean ± SEM with individual data points overlaid. Statistical significance was assessed by mixed-effects analysis with Sidak's multiple-comparisons test. \* $p < 0.05$, \*\*\* $p < 0.001$, \*\*\*\* $p < 0.0001$.

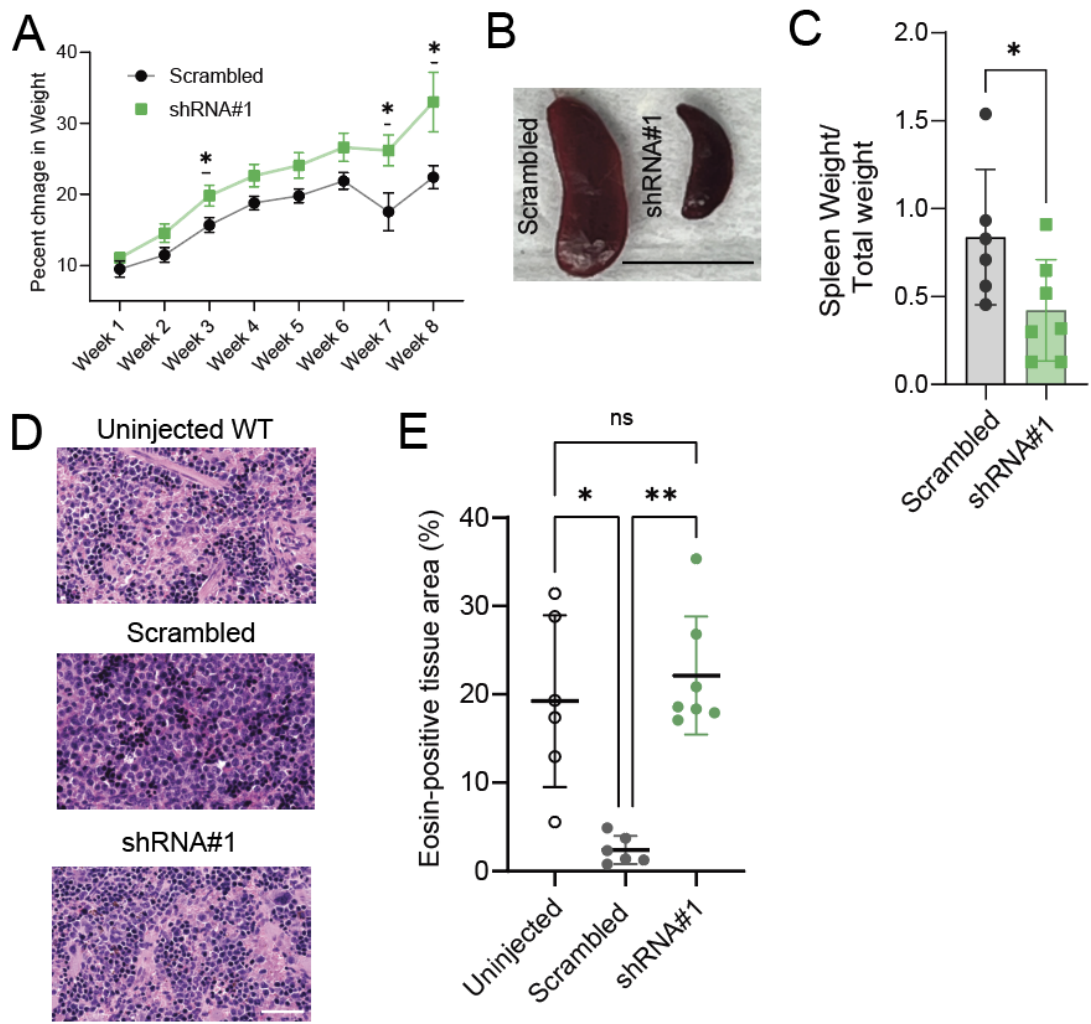

**Supplemental Figure 8. NHE1 knockdown is associated with reduced splenic involvement in T-ALL xenografts.** **(A)** Percent change in body weight over 8 weeks in immunodeficient mice transplanted with luciferase-expressing Jurkat cells expressing scrambled control or NHE1-targeting shRNA#1. Data are shown as mean ± SEM. Statistical significance was assessed using multiple unpaired t-tests for each week **(B)** Representative gross spleen images from mice transplanted with scrambled control or shRNA#1 cells. Scale bar, 1 cm. **(C)** Spleen weight normalized to total body weight in mice transplanted with scrambled control or shRNA#1 cells. Each point represents one mouse; data are shown as mean ± SD. Statistical significance was assessed using a two-tailed Mann–Whitney test. **(D)** Representative hematoxylin and eosin (H&E)-stained spleen sections from uninjected wild-type (WT) mice and mice transplanted with scrambled control or shRNA#1 cells. Scale bars, 50 μm. **(E)** HALO-based quantification of eosin-positive tissue area in H&E-stained spleen sections from uninjected WT mice and mice transplanted with scrambled control or shRNA#1 cells. Each point represents one mouse; data are shown as mean ± SD. Statistical significance was assessed using a Kruskal–Wallis test with Dunn’s multiple-comparisons test. \* $p < 0.05$; \*\* $p < 0.01$; ns, not significant.

**Supplemental Table 2. Limiting-dilution transplantation data and ELDA estimates for top candidate compounds**

| Group | Cell Dose | Sample Size | Positive | LPC Frequency (95% CI) | P Value |
| --- | --- | --- | --- | --- | --- |
| DMSO | 1000 | 5 | 5 | 1 in 10.2<br>(5.43 - 19.7) | NA |
|  | 100 | 7 | 7 |  |  |
|  | 10 | 14 | 9 |  |  |
| Amiloride | 1000 | 5 | 4 | 1 in 149.6<br>(60.82 – 369.1) | < 0.0001 |
|  | 100 | 8 | 7 |  |  |
|  | 10 | 10 | 3 |  |  |
| Deferoxamine | 1000 | 5 | 5 | 1 in 9.61<br>(4.92 – 19.3) | 0.899 |
|  | 100 | 7 | 7 |  |  |
|  | 10 | 12 | 8 |  |  |
| Oxcarbazepine | 1000 | 4 | 4 | 1 in 43.9<br>(18.81 – 103.5) | 0.004 |
|  | 100 | 7 | 5 |  |  |
|  | 10 | 5 | 4 |  |  |
| Rifampicin | 1000 | 5 | 5 | 1 in 5.07<br>(2.51 – 11.1) | 0.171 |
|  | 100 | 8 | 8 |  |  |
|  | 10 | 9 | 8 |  |  |

**Supplemental Table 3. shRNA sequence used for NHE1 knockdown**

|  | Targeted sequence |
| --- | --- |
| NHE1 shRNA#1 | CCAATCTTAGTTTCTAACCAA |
| NHE1 shRNA#2 | CGGTGAGCAGATCAACAACAT |
| NHE1 shRNA#3 | CATCGGATCTTCCCTTCCTTA |
| NHE1 shRNA#4 | GACAAGCTCAACCGGTTTAAT |

**Supplemental Table 4. Primer sequences used for RT-qPCR**

| Gene | Direction | Primer |
| --- | --- | --- |
| ABCG2 | Forward | TGTTTTGTGTTTATGATGGTCTGT |
|  | Reverse | GCTGCAAAGCCGTAAATCCAT |
| BCL2 | Forward | GGGGTCATGTGTGTGGAGAG |
|  | Reverse | GTTCCACAAAGGCATCCCAG |
| CD34 | Forward | GCCACCAGAGCTATTCCCAA |
|  | Reverse | ATAAGGGTCTTCGCCCAGCC |
| CD3G | Forward | TGCCCAATGACCAGCTCTAC |
|  | Reverse | AGGAGAACACCTGGACTACTCT |
| CD44 | Forward | ACACAAATGGCTGGTACGTCT |
|  | Reverse | TGTGGTTGAAATGGTGCTGG |
| CD7 | Forward | CTATCACCATGCACCGCCTG |
|  | Reverse | TCCTTGGGACTGTTCCTCTGT |
| KLF4 | Forward | CGAACCCACACAGGTGAGAA |
|  | Reverse | ATGCTCGGTCGCATTTTTGG |
| Lgr5 | Forward | AAGCCTTCAATCCCTGCGTC |
|  | Reverse | CCTGGACGGGGATTTCTGTT |
| NANOG | Forward | AATGGTGTGACGCAGGGATG |
|  | Reverse | TGCACCAGGTCTGAGTGTTC |
| POU5F1 | Forward | CTCGAGAAGGATGTGGTCCG |
|  | Reverse | CCTGAGAAAGGAGACCCAGC |
| PROM1 | Forward | GAGCTAAGGGAAGGGCGG |
|  | Reverse | TTCTGTCTGAGGCTGGCTTG |
| Sox2 | Forward | AGGATAAGTACACGCTGCCC |
|  | Reverse | TAACTGTCCATGCGCTGGTT |
| Eef2 | Forward | GGAGCCAAGTGCTAACATGC |
|  | Reverse | CTCGCCAACAGGAACAGTA |
| RPL19 | Forward | GAGCTCTTTCCTTTCGCTGCT |
|  | Reverse | CCGGATCTGCTGACGGGAG |
| ACTB | Forward | ACAGAGCCTCGCCTTTGC |
|  | Reverse | CCACCATCACGCCCTGC |
| SLC9A1 | Forward | GCCTCATGAAGATAGGTTTCCAT |
|  | Reverse | GTGTCTCGCCTACACCCTTG |
