## Supplemental Methods for "NHE1 sustains leukemia-propagating capacity and mitochondrial fitness in T-cell acute lymphoblastic leukemia"

#### **S1. Zebrafish care and use**

All zebrafish studies were approved by the University of Kentucky Institutional Animal Care and Use Committee (IACUC; protocol 2019-3399). Syngeneic CG1-strain zebrafish were used (1). Fish were maintained at 28°C on a 14-hour light/10-hour dark cycle. Embryos were collected in 1× E3 medium containing 5.0 mM NaCl, 0.17 mM KCl, 0.33 mM CaCl<sub>2</sub>, 0.33 mM MgSO<sub>4</sub>, and 200 µL/L methylene blue. For transplantation or imaging, fish were anesthetized in system water containing 200 µL of a 4 mg/mL MS-222 stock solution per 25 mL.

#### **S2. Propagation of zebrafish T-ALL clones**

Characterization of zebrafish T-ALL clones 1.3 and 10.2, enriched for leukemia propagating cells, has been previously described (2). Frozen cells were thawed and expanded as previously described (3). Briefly, vials frozen cells were rapidly thawed, cells were washed in 10mL zebrafish cell medium (0.9× PBS with 5% FBS), and centrifuged at 2,500 × g at 4°C. Pellets were resuspended in 0.5 mL cell medium and kept on ice. For propagation, approximately 50,000 cells per 5 µL were prepared and injected intraperitoneally into CG1 recipients using a 26-gauge, 2-inch Hamilton microsyringe. Animals were monitored weekly for leukemia progression.

#### **S3. Primary in vivo drug screen**

Zebrafish T-ALL clones 1.3 and 10.2 were propagated in CG1 zebrafish. After approximately three weeks, fish with approximately 80% leukemia burden were euthanized. Leukemia cells were recovered by macerating the fish and passing the suspension through a 40 µm cell strainer. Fluorescent leukemia cells were counted by microscopy, and viability was assessed using Trypan Blue staining. The Enzo Screen-Well FDA Approved Drug Library V2 (BML-2843-0100, version 1.4, Lot No. 06051910A) was used. Most compounds were supplied in DMSO, with a minority in water, as a stock concentration of 10mM. Compounds were pooled by plate column (8 compounds per pool) and diluted in RPMI 1640 + supplemented with 10% FBS media to 1 µM per compound. Drug-pool identifiers corresponded to plate and column positions (for example, P2C5 refers to the drug group pooled plate 2, column 5).

Leukemic cells were exposed to each drug pool for 1 hour at room temperature before the drug-cell suspension was transplanted intraperitoneally into syngeneic CG1 zebrafish (15 recipients per treatment) at a dose of 25 leukemia cells per fish. Fish were monitored weekly until all animals in the control group developed leukemia, which was the primary screen endpoint. Drug-pool activity was expressed relative to the corresponding vehicle treated leukemia clone. Pools meeting the prespecified leukemia-incidence and animal viability criteria were advanced to secondary screening.

#### **S4. Limiting dilution transplantation into adult CG1 zebrafish**

Limiting dilution transplantation was used to quantify LPC frequency, as previously described (3–5). Briefly, fish with ~80% leukemia burden (clone # 1.3) were recovered, filtered, and counted. Cells were diluted in RPMI 1640 + 10% FBS to 10, 100, or 1,000 cells per 5 µl and incubated for 1 hour with 1 µM of compound or the corresponding vehicle control. Each cell solution/drug mixture was transplanted intraperitoneally into CG1 recipients using a Hamilton micro-syringe. Fish were housed in 4-L tanks and examined weekly for fluorescent leukemia development. At 6 weeks, recipients with any detectable GFP-positive leukemia signal were scored as positive. LPC frequency and 95% confidence intervals were estimated with the web-based ELDA (Extreme Limiting Dilution Analysis) statistical software (6).

### **S5. Human T-ALL cell culture and drug treatments**

Jurkat (ATCC TIB-152), CCRF-CEM (ATCC CCL-119), and MOLT-4 (ATCC CRL-1582) cells were cultured in RPMI 1640 (Thermo Fisher Scientific, 11875119) containing 10% heat-inactivated FBS (Atlanta Biologicals, S11150H) and 1× penicillin–streptomycin–glutamine (Thermo Fisher Scientific, 10378016) at 37°C and 5% CO<sub>2</sub>. Cell cultures were screened routinely for mycoplasma using the LookOut Mycoplasma PCR Detection Kit (Sigma, MP0035-1KT). Cells were used between passages 4 and 20 after thawing.

For pharmacologic studies, cells with >90% viability were treated with amiloride, hexamethylene amiloride (HMA), or the indicated screening compounds and vehicle controls in standard culture media. Compound concentrations, incubation times, and assay-specific treatment conditions are specified in the figure legends.

### **S6. Lentiviral NHE1 knockdown**

HEK293T cells were transfected with psPAX2, pCMV-VSV-G, and the shRNA construct of interest using TransIT-LT1 transfection reagent (Mirus Bio, MR2300), following the manufacturer's instructions. psPAX2 was a gift from Didier Trono (Addgene plasmid # 12260 ; http://n2t.net/addgene:12260 ; RRID:Addgene\_12260), pCMV-VSV-G was a gift from Bob Weinberg (Addgene plasmid # 8454 ; http://n2t.net/addgene:8454 ; RRID:Addgene\_8454) (7). A scrambled sequence in the pLKO.1 backbone was used as a control. scramble shRNA was a gift from David Sabatini (Addgene plasmid # 1864 ; http://n2t.net/addgene:1864 ; RRID:Addgene\_1864) (8). NHE1/SLC9A1-targeting shRNAs vectors were purchased from Sigma; shRNA sequences are provided in **Supplementary Table 2**. Viral supernatants were collected at 48 and 72 hours after transfection.

For transduction, 2.5 mL of viral supernatant was added to $0.5 \times 10^6$ Jurkat cells with 10 µg/mL polybrene (ThermoFisher, TR-1003-G), followed by centrifugation at 2,250 rpm for 1.5 hours in a six-well plates. After 24 hours, cells were washed twice with 1X PBS and selected with 10 µg/mL puromycin for 1 week before use. NHE1 knockdown was verified by immunoblotting and RT-qPCR.

### **S7. Colony formation assay**

Colony formation was assessed as previously described (3,9). Viable cells (>90% viability) were treated with drugs or controls where applicable and mixed with MethoCult™ H4100 base medium (Stem Cell Technologies, 04100) supplemented with 20% FBS. Suspensions were plated in 24-well plates at 2,000 cells per well in 0.5 ml. Colonies were imaged 14 days later at 4x magnification using an Agilent BioTek Lionheart FX instrument. Images were stitched using Gen5 software, and colonies were counted in ImageJ.

### **S8. Apoptosis and cell cycle analysis**

For apoptosis measurements, cells were treated at the indicated drug concentrations and times where applicable, collected, and stained with the Annexin V-FITC kit (Invitrogen, V13242A), according to the manufacturer's protocol, with propidium iodide (ThermoFisher, P1304MP) used where indicated. Samples were acquired on a BD FACSymphony A3 flow cytometer and analyzed using FlowJo.

For cell cycle analysis, cells were washed in 1X PBS and fixed in ice-cold 70% ethanol for at least 30 minutes. Fixed cells were washed with 1X PBS and incubated with PureLink RNase (100 µg/mL; Invitrogen 12091-021) and propidium iodide (30 µg/mL) in 1X PBS for 15 minutes at room temperature in the dark. Samples were filtered through a 40 µm cell strainers, analyzed on a BD Symphony A3, and modeled in FlowJo.

### **S9. Immunoblotting and RT-qPCR**

Cells were lysed in M-PER buffer (ThermoFisher, 78503) containing protease inhibitors and EDTA (Boster, AR1182). Healthy donor PBMCs lysates used for comparison were obtained from Stem Cell Technologies (70025.1). Primary antibodies included NHE1 (1:1000; Santa Cruz Biotechnology, sc-136239), GAPDH (1:1000; Cell Signaling Technology, 2118L), and Tubulin (1:1000; Cell Signaling Technology, 2148S). Blots were blocked in 5% milk in 0.1% TBST for 1 hour, incubated with primary antibodies overnight at 4°C, washed three times with 0.1% TBST, and incubated for 1 hour at room temperature with species-appropriate HRP-conjugated secondary antibodies for 1 hour. Secondary antibodies included anti-mouse IgG (1:1000; Cell Signaling Technology, 7076P2) and anti-rabbit IgG (1:1000; R&D systems, HAF008). Bands were visualized using the Clarity Western ECL Substrate (Bio-Rad, 1705060) on a ChemiDoc Touch system and quantified using ImageJ.

RNA isolation and real-time qRT-PCR was done as previously described (3). Primer sequences are listed in **Supplemental Table 3**. Gene expression was normalized to reference gene expression using the $2^{-\Delta\Delta Ct}$ method.

### **S10. Seahorse mitochondrial stress testing**

Cells with >90% viability were seeded at 60,000 – 100,000 cells per well (optimized for each cell line) in Cell-Tak (Corning, 354240) coated Agilent Seahorse XF Pro 96-well plates. After carefully pipetting the cells into the wells, plates were centrifuged at $200 \times g$ for 1 minute without braking, then incubated in a 37 °C incubator without CO<sub>2</sub> for 25–30 minutes to promote adherence. Measurements were performed in serum-free, phenol red-free RPMI assay media.

Oxygen consumption was measured over approximately 90 minutes following sequential injection of oligomycin A (1 µM), FCCP (0.6–2 µM, optimized for cell line), and rotenone plus antimycin A (1 µM each). Basal respiration, ATP-linked respiration, and spare respiratory capacity were calculated using Seahorse Wave software (version 2.6.3). Data were normalized to cell number or total protein per well, as appropriate for each experiment. Drug exposure conditions are reported in the figure legends.

### **S11. Confocal mitochondrial imaging and morphology analysis**

For mitochondrial visualization and morphology analysis, 100,000 cells per well were seeded in 96-well black glass-bottomed plates (Cellvis, P96-1.5H-N) and allowed to settle or adhere for 30 minutes (10). Cells were then fixed in 4% paraformaldehyde (Electron Microscopy Sciences, 157-4-100) for 10 minutes, washed in 1X PBS rinse, and stained with Hoechst (Invitrogen, H3570; 1:1000), Rhodamine Phalloidin (Abcam, ab235138; 1:500), and MitoView™ Green (Biotium, 70054; 1:200) for 1 hour at room temperature. Cells were rinsed with 1X PBS prior to imaging. Images were acquired in the University of Kentucky Light Microscopy Core using a Nikon A1R confocal microscope and 60X oil-immersion objective. Image overlays and display adjustments were prepared in Adobe Photoshop 2020. Unprocessed Nikon .nd2 files were converted to .tif file and analyzed in ImageJ using the Mitochondrial Network Analysis (MiNA) plugin (11).

### **S12. Flow cytometry for mitochondrial phenotypes**

Mitochondrial content was assessed using by MitoView™ Green (Biotium, 70054), and mitochondrial membrane potential using JC-1 staining (Biotium, 30001). Staining was performed according to the manufacturers' instructions, using appropriate controls. Cells were filtered through a 40

µm cell strainer (VWR, 21008-948) before acquisition on a BD Symphony A3. Data were analyzed using FlowJo.

#### **S13. RNA sequencing and pathway analysis**

For transcriptomic profiling, RNA was isolated using the Quick-RNA kit (Zymo Research, R1054). Library preparation was performed using the NEBNext Ultra II Express Library Prep Kit with the Poly(A) mRNA Magnetic Isolation Module, following the manufacturer's instructions and as previously described (12). Libraries were assessed with an Agilent Bioanalyzer, pooled equimolarly, and sequenced with 2 × 150-bp paired-end reads on an Illumina NovaSeq sequencing platform. FASTQ files were processed in Partek Flow version 12.3. Reads were aligned to the human reference genome with STAR and quantified with HTSeq. Differential gene expression was analyzed using DESeq2 with gene set enrichment analysis (GSEA) performed using the GSEA software (13). RNA-seq data were deposited in the NCBI Gene Expression Omnibus under accession GSE341718.

#### **S14. Proteomic profiling and pathway analysis**

Protein samples were prepared with the PreOmics iST kit. Samples were lysed in 100 µL of lysis buffer, heated to 95°C for 10 minutes, and digested at 37°C for 3 hours. Peptides were vacuum-dried and resuspend in 5% acetonitrile with 0.1% formic acid. Analysis was performed with a Thermo Scientific™ Orbitrap Exploris 240 mass spectrometer and a Vanquish Neo UHPLC system using an EASY-Spray HPLC. Peptides were separated using a linear gradient from 2-55% mobile phase B over 90 minutes at 0.3 µL/min. Data-dependent acquisition used a 60k MS1 resolution, scan range of 375-1200 m/z and 15k MS2 resolution with 2 m/z isolation width, and 26% normalized HCD collision energy. Raw spectra were processed in Proteome Discoverer version 3.1 with the SEQUEST algorithm against the Homo Sapiens UniProt database (downloaded March 2024). Abundances were normalized to total peptide abundance.

Differential protein abundance analysis used an adapted workflow based on the DEP R package (14). Pathway enrichment was performed using the WebGestalt (15). Proteins abundance data are presented in **Supplementary Table 4**.

#### **S15. Gas-chromatography spectrometry metabolomics**

Cells were washed twice with ice-cold 0.9% NaCl, and metabolites were extracted using 1 ml of 80% methanol at -20°C containing 20 µM shikimic acid as an internal standard. Samples were held at -20°C for 1 hour and centrifuged at 17,000 x g for 10 minutes at 4°C. Supernatants were transferred to fresh 1.5 ml polypropylene tubes. The insoluble material (protein) was rinsed twice with 50% methanol and once with 100% methanol, dried, resuspended in 2% SDS and 62.5 mM Tris pH 6.8, and denatured by heating at 95°C for 10 minutes. Protein concentration was determined using the bicinchoninic acid assay.

For GC-MS derivatization, extracts were evaporated using a Centrivap system and derivatized with 50 µl of 20 mg/ml methoxyamine in pyridine at 30°C for 90 minutes. Samples were centrifuged at 17,000 x g for 10 minutes at room temperature, after which 40 µl was transferred to a glass GC-MS vial and combined with 80 µl of N-Methyl-N-(trimethylsilyl)trifluoroacetamide (MSTFA) containing 1% chlorotrimethylsilane (TCMS). Sample were incubated at 37°C for 30 minutes and held overnight at room temperature before analysis.

Metabolites were analyzed on an Agilent 5977B GC-MS equipped with an HP-5MS column. The sample injection volume was 1 µL. The inlet temperature was 250°C with 4.9 psi pressure and 3 mL/min septum purge; the splitless inlet was set to 10.5 mL/min at 1 minute. The oven program began at 60°C for 1 minute, followed by a linear ramp to 325°C at a rate of 10°C per minute. Full-scan spectra were collected across 50-550 m/z, with source and quadrupole temperatures of 230°C and 150°C, respectively. Peaks were extracted in Agilent MassHunter Qualitative Analysis. Identified metabolite peak areas were normalized to the peak area of the shikimic acid internal standard and total protein. Data were analyzed in MetaboAnalyst (16). Metabolites abundance data are available in **Supplementary Table 5**.

### **S16. Analysis of public TARGET-ALL P2 data**

TPM-normalized RNA-sequencing data from 463 primary and 69 partially paired recurrent acute lymphoblastic leukemia (ALL) samples in the TARGET-ALL-P2 cohort were obtained from the Genomic Data Commons (GDC) Data Portal (https://portal.gdc.cancer.gov/). A linear mixed-effects model with a random intercept was used to compare *SLC9A1* (NHE1) expression between primary and recurrent leukemia samples, accounting for the partially paired study design.

Processed survival data from the TARGET-ALL-P2 cohort were obtained from cBioPortal (https://www.cbioportal.org/). Kaplan-Meier survival analysis and log-rank tests were used to compare outcomes between patients with high and low *SLC9A1* expression. Expression thresholds evaluated included the median, 25th and 75th percentiles, and an optimized cutoff determined using the R package *maxstat*. For analyses using the optimized cutoff, log-rank *p* values were adjusted for multiple comparisons. Hazard ratios were estimated using Cox proportional hazards regression.

### **S17. NSG mouse xenografts and bioluminescence imaging**

All mouse studies were approved by the University of Kentucky IACUC (protocol 2017-2754). NSG mice (NOD.Cg-Prkdcscid Il2rgtm1Wjl/SzJ; Jackson Laboratory) were used. Jurkat cells were transduced with a luciferase expression vector (Addgene 21471), selected with 600 µg/mL neomycin, and subsequently transduced with plasmids containing either the scrambled or NHE1-targeting shRNA. Male and female mice aged 6 and 8 weeks were randomized to receive either control (n=6) or NHE1 knockdown (n=7) cells. Cells were injected intravenously in 100 µL 1X PBS. Within each group, half of the mice were injected with 10<sup>6</sup> cells and the other half with 10<sup>5</sup> cells to allow stratification of leukemia onset. Animals were followed for up to 10 weeks.

For bioluminescence imaging, mice received 200 µL of D-luciferin (25 mg/mL; G-Biosciences, RC-231). After 7 minutes, animals were anesthetized with isoflurane in a preconditioned chamber and imaged using a Lago SII imaging station (Spectral Instruments Imaging) at a 60-second exposure. Images were quantified using Aura software.

### **S18. Marrow CD45 flow cytometry**

Femurs and tibiae were isolated after euthanasia and cleaned of surrounding tissue using a Kimwipe soaked in 70% ethanol. Both ends of each bone were cut with sharp scissors, and marrow was flushed into RPMI medium supplemented with 10% FBS using a 25-gauge needle. Suspensions were filtered through 40-µm strainers and washed twice in 1X PBS. Cells were counted and stained with PE-conjugated anti-human CD45 antibody (BioLegend, 304008; 5 µL per 10<sup>6</sup> cells in 100 µL staining volume). DAPI (Thermo Fisher Scientific, 62248; 1:1,000) was used for exclusion of nonviable cells. Cells were analyzed on a before acquisition on a BD Symphony A3. Data were analyzed using FlowJo.

### **S19. Spleen histology and marrow cleaved caspase-3 staining**

Spleens were harvested at euthanasia, fixed overnight in 10% neutral-buffered formalin with gentle agitation, then transferred to 70% ethanol and stored at 4°C. Spleens were paraffin embedded in paraffin, sectioned, and stained with Hematoxylin and Eosin (H&E). Femurs for marrow histology were similarly fixed, decalcified using OSTEOMOLL (Sigma, 12352401), paraffin embedded, sectioned at 4 µm and mounted on positively charged slides before baking at 58°C for a minimum of 1 hour.

Cleaved caspase-3 staining was performed on a Ventana Discovery Ultra Autostainer using CC1 antigen retrieval and anti-cleaved caspase-3 (Cell Signaling Technology, 9661; 1:150) for 1 hour, followed by OmniMap anti-rabbit-HRP (Roche, 760-149) and ChromoMap DAB (Roche, 760-159), following the manufacturer's recommendations. Slides were counterstained with Mayer's hematoxylin and blued with ammonia water prior, dehydrated, cleared, and mounted.

Slides were scanned at 20× or 40× magnification using an Aperio slide scanner and analyzed using HALO (Indica Labs). Cleaved caspase-3 positivity was quantified by assessing nuclear color intensity with the Multiplex IHC algorithm (version 3.4.9). Histological and staining were performed by the Markey Cancer Center Biospecimens Procurement and Translational Pathology Shared Resource Facility.

### **S20. Statistical analyses**

Data visualization and most statistical analyses were conducted using GraphPad Prism 10.4.1. Depending on the experimental design, statistical tests included two-tailed Student's or Welch's t-tests, one- or two-way ANOVA with appropriate multiple-comparison correction (including Dunnett's test), Gehan–Breslow–Wilcoxon tests, and log-rank (Mantel–Cox) analyses. ELDA was used for zebrafish leukemia-propagating cell-frequency estimates. Clinical survival analyses also included Cox proportional hazards modeling and the maxstat cutoff procedure. Unless otherwise indicated, bar graph error bars represent standard deviations. Cell-based experiments were performed with at least three biological replicates. Statistical significance was defined as $p < 0.05$. Asterisks denote \*$p < 0.05$, \*\*$p < 0.01$, \*\*\*$p < 0.001$, and \*\*\*\*$p < 0.0001$. Sample sizes, specific tests, and multiple-testing adjustments are provided in the corresponding figure legends.
